# Genomic programming of antigen cross-presentation in IRF4-expressing human Langerhans cells

**DOI:** 10.1101/541383

**Authors:** Marta E Polak, Sofia Sirvent, Kalum Clayton, James Davies, Andres F. Vallejo, Zhiguo Wu, Jeongmin Woo, Jeremy Riddell, Patrick Stumpf, Matthew Rose-Zerilli, Jonathan West, Mario Pujato, Xiaoting Chen, Christopher H. Woelk, Ben MacArthur, Michael Ardern-Jones, Peter S Friedmann, Matthew T. Weirauch, Harinder Singh

**Affiliations:** Clinical and Experimental Sciences, Sir Henry Wellcome Laboratories, Faculty of Medicine, University of Southampton, SO16 6YD, Southampton, UK; Institute for Life Sciences, University of Southampton, SO17 1BJ, UK; Division of Immunobiology & Center for Systems Immunology, Cincinnati Children’s Hospital Medical Center, Cincinnati, OH, 45229, USA; College of Life Sciences, Wuhan University, Wuhan 430079, Hubei Province, China; Samsung Genome Institute, Samsung Medical Center, Seoul, South Korea; Center for Autoimmune Genomics and Etiology, Cincinnati Children’s Hospital Medical Center, Cincinnati, OH, 45229, USA; Centre for Human Development, Stem Cells and Regeneration, Faculty of Medicine, University of Southampton, Southampton SO17 1BJ, UK; Cancer Sciences, Faculty of Medicine, University of Southampton, SO16 6YD, Southampton, UK; Merck’s Exploratory Science Center, Cambridge, MA 02141, USA; Department of Pediatrics, University of Cincinnati College of Medicine, Cincinnati, Ohio, 45229, USA

**Keywords:** Langerhans cell, Interferon regulatory factor 4, Interferon regulatory factor 8, antigen cross-presentation, human skin, transcriptional programming, chromatin landscape, gene regulation

## Abstract

Langerhans cells (LCs) in the epidermis present MHC I and MHC II-restricted antigens thereby priming either CD8 or CD4 T cell immune responses. The genomic programs and transcription factors regulating antigen presentation in LCs remain to be elucidated. We show human LCs are highly efficient in MHC I-antigen cross-presentation but lack the transcription factor IRF8 that is critical in dendritic cells. LC migration from the epidermis enhances their ability to cross-present antigens and is accompanied by the induction of the transcription factor IRF4, whose expression is correlated by scRNA-seq with genes involved in ubiquitin-dependent protein degradation. Chromatin profiling reveals enrichment of EICE and AICE composite DNA binding motifs in regulatory regions of antigen-presentation genes, which can be recognized by IRF4 in conjunction with PU.1 or BATF3 expressed in LCs. Thus, the genomic programming of human LCs including inducible expression of IRF4 with enhanced cross-presentation distinguishes them from conventional dendritic cells.

## Introduction

Langerhans cells (LCs) reside in the epidermis as a dense network of immune system sentinels. They are uniquely specialised at “sensing” the environment by extending dendrites through inter-cellular tight junctions to gain access to the *stratum corneum*, the outermost part of the skin. LC are highly specialised antigen presenting cells, priming protective immune responses against pathogens encountered via the skin, such as viruses ^1–3^, bacteria ^4^ and fungi ^5^. They also promote responses to chemical sensitizers ^6,7^. Their position in the outermost layers of the skin barrier and their capacity to “sense” dangerous perturbations to their environment make them a critical first line of defence in the skin.

LCs were first shown to be highly efficient at presenting exogenous antigens in the context of MHC Class II thereby priming antigen-specific CD4 T cells ^8^. More recently, they have also been shown to be capable of efficient cross-presentation in which exogenous antigens are presented on MHC class I, resulting in activation and expansion of antigen-specific effector CD8 T cells ^2,5,9^. Such cross-presentation becomes particularly important for adaptive immune responses against viruses and also cancerous cells which have evolved immune evasion mechanisms that inactivate DCs ^10,11^.

Activation of skin immune responses requires participation of epidermal cells in collaboration with LCs; cross-talk between the epidermal and immune components being critical. For example, in models of cutaneous viral infection, including vaccinia virus, only skin structural cells support virus replication, while immune cells, including LCs, are infected abortively ^12^, necessitating antigen transfer between structural and immune skin components. Cytokine signalling from keratinocytes has been shown to impact LC development and function. Epidermal TGF-β and BMP7 are required for LC development and tolerogenic function *in situ* ^13–15^ and cytokine signalling via thymic stromal lymphopoietin (TSLP) in atopic skin disease polarises skin dendritic cells to prime Th2 and Th22 CD4 T cell responses, while reducing the ability of LCs to cross-prime CD8 T cells ^16–19^. In contrast, TNF-α, a pro-inflammatory cytokine released in skin by a variety of cell types including keratinocytes and fibroblasts as well as dermal macrophages and neutrophils, provides a key component of cutaneous anti-viral immune responses. Numerous reports demonstrate, that keratinocyte-derived TNF-α delivers highly potent signals inducing LC immunogenic function and ability to present antigens ^2,20–23^ and enhances their ability to prime antiviral adaptive immunity ^24^. Cross-talk between LC and surrounding keratinocytes, coupled with their ability to cross-present antigens expressed by other cells to skin resident and infiltrating T lymphocytes, defines the major role of LC in skin immunity and allows them to initiate efficient adaptive immune responses in the context of skin infection ^25–27^. However, little is known about the genomic mechanisms which programme LC functions in homeostasis and inflammation and how epidermal-derived signals modify such programming.

Like macrophages, LCs originate from yolk-sac progenitors and populate the epidermis during embryonic life. However, functionally, they are more similar to conventional DCs in their ability to efficiently present and cross-present antigens to prime T cell responses ^28,29^. Our recent research suggests that even though LCs are among the most efficient antigen presenting cells, their transcriptional networks are distinct from those of both macrophages and DCs ^22,30^, warranting deeper analyses. Molecular studies and comparative transcriptome analysis further highlight the fundamental differences in gene regulatory networks between LCs and conventional DCs ^30,31^. In recent years, in all DC subtypes studied, two members of the interferon regulatory factor (IRF) family, IRF-4 and IRF-8, have emerged as key players in DC development and function ^32–35^. IRF4 and IRF8 are critical for all aspects of DC biology, controlling a wide range of functions. These include induction of innate responses elicited via pattern recognition receptors TLR, NOD, and RIG during viral and bacterial infection ^36–38^, cell activation, and efficient antigen uptake, presentation and cross-presentation ^23,33,39^, migration and tolerance induction ^39,40^. Interestingly, it has been postulated that the ability of murine DC subsets to efficiently activate CD8 and CD4 T cells, depending on the presentation of antigen in the context of MHC class I and II, is determined by the relative expression of IRF8 and IRF4, respectively ^33^. Furthermore, the function of conventional DCs, excelling at cross-presentation of tumor cell-associated antigens, is critically dependent on BATF3/IRF8 complexes ^41,42^. Genome-wide expression analysis reveals similarities between human LCs and cross-priming murine dendritic cells ^29,30^. LC development has been shown to be IRF4 and IRF8-independent in both murine and human systems ^31,43,44^. Crucially, it remains to be determined if IRF4 and/or IRF8 regulate the genomic programming and functions of human epidermal LCs.

Here we show that human LCs enhance their ability to cross-present antigens during their migration out of the epidermis. This is accompanied by the induction of the transcription factor IRF4 in the absence of IRF8. Furthermore, expression of IRF4 in human LCs is correlated by scRNA-seq with genes involved in ubiquitin-dependent protein degradation. Priming of CD8 T cell responses by LCs is further stimulated by TNF-α signalling. Chromatin profiling before and after TNF-α stimulation reveals enrichment of EICE and AICE transcription factor binding motifs in regulatory regions of antigen-presentation gene modules. These composite motifs can be recognized by IRF4 in conjunction with PU.1 and BATF3, respectively, also expressed in these cells. Thus, the genomic programming of LCs for cross-presentation is independent of IRF8 and instead appears to utilize IRF4 in combination with PU.1 and BATF3, thereby differentiating LC from conventional dendritic cells.

## Results

### Migration and TNF-α signalling induce antigen cross-presentation and priming of CD8 T cell responses by human skin LCs

To enable the analysis of the genomic programming of primary human LCs, we utilized established protocols for isolating highly pure populations of viable and functional LCs from the epidermis, (^2,22^, Figure 1a-c, Supplementary data Figure S1). In agreement with our earlier findings and those of others ^2,9,45^, human LC after migrating from the epidermis uniformly expressed high levels of CD1a, CD207 and HLA-DR (Figure 1b, Supplementary data Figure S1). Such migrated LCs are able to take up and present antigens to CD8 and CD4 T cells, activating the latter (^2,23^, Supplementary data Figure S1). These LCs express high levels of MHC class I and II complexes on their surface. Notably, migration from the epidermis increases LC activation status, as assessed by enhanced expression of co-stimulatory molecules: CD40, CD70, CD80, CD86 (Figure 1c). To analyse the effects of migration as well as TNF-α signalling on LC cross-priming we used an established model for antigen cross-presentation ^2^. Steady state and migrated LCs were pulsed with a 30 amino acid peptide, comprising a 9 amino acid HLA-A2 restricted GLC epitope from Epstein Barr Virus protein BMLF and stimulated with TNF-α. We have previously demonstrated that the cross-presentation of the GLC epitope to antigen-specific CD8 T cells was critically dependent on the ability of LCs to take up and actively process the 30AA peptide for presentation ^2^. LC migration from the skin induced their ability to cross-present antigens as measured by IFN-γ release from a GLC peptide-specific HLA-A2 CD8 T cell line (Figure 1d, Supplementary data Figure S1). TNF-α signalling further enhanced the ability of migrated LCs to cross-present the same antigen (Figure 1e). We note that in the presence or absence of TNF-α, cells fixed with glutaraldehyde prior to antigen pulsing did not activate cognate CD8 T cells. In contrast, fixation did not affect presentation of a short peptide, which could be externally loaded onto the MHC molecules. Furthermore, fixing LC post pulsing but before co-culture with T lymphocytes, reduced their ability to activate CD8 T cells ^2^. This is consistent with the inhibition of intracellular protein trafficking and antigen processing by glutaraldehye fixation. Thus, LC migration upregulates co-stimulatory molecules and their antigen cross-presentation capabilities, the latter can be further augmented by TNF-α signalling.

**Figure 1.**
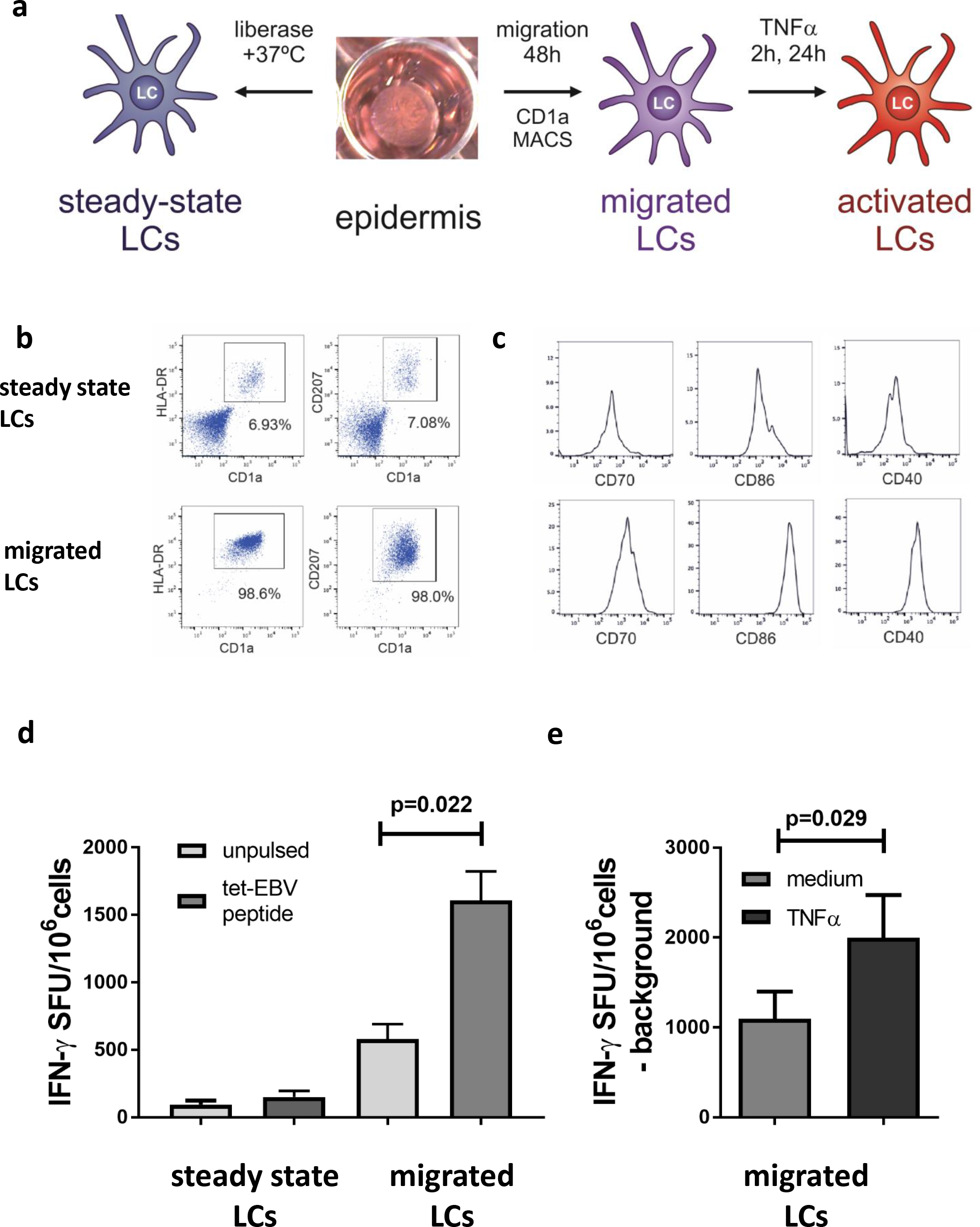
Migration and TNF-α signalling induce antigen cross-presentation by human skin LCs and priming of CD8 T cell responses. a. Schematic depicting isolation of primary human LCs. Split healthy skin was treated with dispase for 20h to dissociate epidermis. Steady-state LCs were isolated from the epidermis by digestion with liberase TM or migrated from the epidermal sheets for 48h in cell culture medium and stimulated with TNF-α to induce their activation.
b. Flow cytometry assessment of steady-state and migrated LC. LCs were enumerated as CD207/CD1a/HLA-DR high cells. A representative example of n>5 experiments.
c. Flow cytometry assessment of activation markers expressed by steady-state and migrated LC. Co-stimulatory molecules critical for CD8 T cell activation during antigen presentation (CD70, CD86 and CD40) were analyzed in CD207/CD1a/HLA-DR high cells. A representative example of n>5 experiments.
d. IFN-γ secretion by an EBV-specific CD8 T cell line stimulated with antigen presenting LCs in the context of MHC I HLA-A2. Steady-state or migrated LCs were pulsed with 30 amino acid peptide containing EBV epitope (dark grey). Pulsed or unpulsed (light gray) LCs were stimulated with TNF-α and then assayed for IFN-γ secretion. ELISpot assay, n=5 independent experiments in triplicate.
e. IFN-γ secretion by EBV-specific CD8 T cell line stimulated by migrated LCs pulsed as in panel 1d. IFN-γ secretion was measured with (black) or without (grey) TNF-α stimulation. ELISpot assay, n=5 independent experiments in triplicate

### Human skin LCs are transcriptionally programmed to efficiently cross-present antigens

To gain insights into the genomic programming of migrated LCs we analysed their transcriptome using bulk RNA-sequencing. The antigen processing and presentation genes were quantitatively amongst the highest expressed genes in migrated LCs and are therefore designated as the core LC transcriptional programme (Figure 2a, Supplementary data Figure S2). This confirmed and extended our previous analysis using DNA microarrays ^22^. We next compared the expression of genes in migrated LCs with previously reported signatures of cross-presenting DCs, including those in the Reactome database and reported by Artyomov and colleagues ^29^. These were compiled into an “antigen cross-presenting” molecular signature (Supplementary data Table 1, Supplementary Figure S2) and “antigen processing” signature (Figure 2b). In agreement with previously published data ^29^, the gene signature encoding antigen processing and presentation in different populations of cross-presenting human skin and blood derived DCs was recapitulated in human LCs, suggesting the existence of a shared transcriptional programme (Figure 2b, Supplementary data, Figure S2). While 57 genes shared between all three subsets encoded for proteasome structure (41 genes, FDR p= 7.32^−93^), protein catabolic process, (FDR p = 8.164^−100^) and antigen presentation to class I (FDR p = 1.324^−95^), 287 genes shared between LCs and other cross-presenting DCs were involved in metabolic processes (FDR p = 5.39^−19^), including NADH dehydrogenase activity FDR p = 2.723^−11^). Notably, the core LC genomic programme was accompanied by high levels of expression of genes encoding ubiquitin protease activity (Figure 2a). Accordingly, 64/66 genes shared between LCs and the antigen processing signature encoded protein ubiquitination components (FDR p = 1.340^−83^).

**Figure 2.**
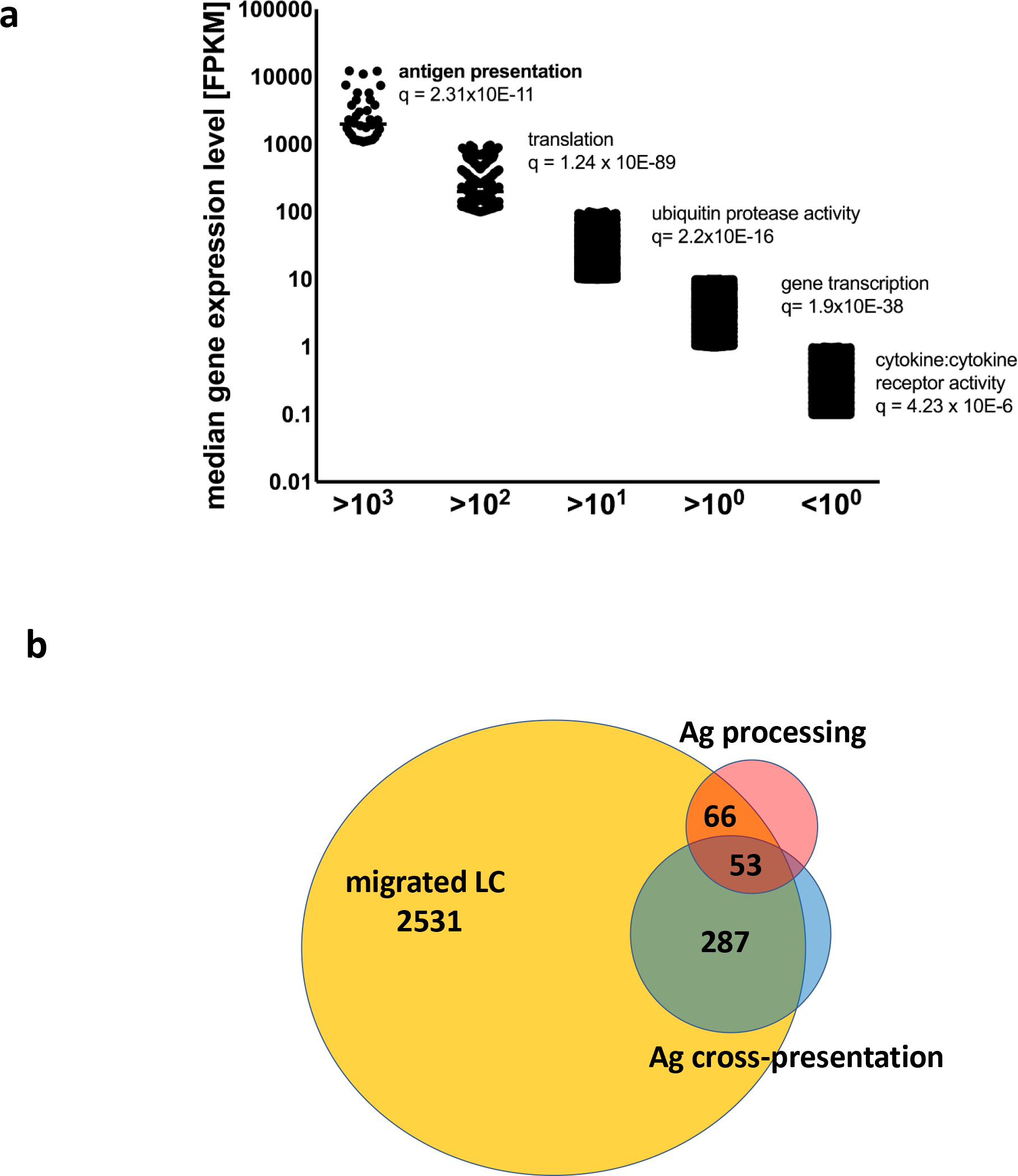
Human skin migrated LCs are programmed at the transcriptome level to efficiently cross-present antigens. a. Dominant biological processes and pathways enriched in genes expressed at varying levels in steady state migrated LCs. Gene ontology analysis for each expression level (FPKM) interval determined by RNA-seq was performed using ToppGene on-line tool ^91^. Top unique Biological processes are shown for each interval, significance denoted by FDR (Benjamini-Hochberg) corrected p-value is shown.
b. Overlaps between reported cross-presentation (373 genes) and antigen processing (212 genes) signatures, and genes expressed in steady-state migrated LC >10 FPKM.

### Single cell transcriptome profiling reveals correlation between metabolism and antigen processing gene modules in human LCs

Next, we used scRNA-seq to analyse the transcriptomes of migrated single cells isolated from human epidermis, a population highly enriched in LCs (>80% of CD1a+, HLADR+ cells, Supplementary data Figure S3). Seurat analysis ^46^ clearly identified two main cell subsets, confirming LC identity for 138 out of 150 cells (92%, p = 1.163^−21^), and assigning the remaining population as melanocytes (p=9.624^−9^) (Supplementary data Figure S3). Interestingly, as identified by Seurat analysis, the main genes defining the LC cluster features were those encoding antigen presenting molecules *HLADR* and *CD74*, together with *TMSB4X*, which is involved in actin polymerisation, cell motility and cytoskeleton re-organisation.

We next mapped single cell LC transcriptomes onto the recently described 10 sub-populations of human blood monocytes and DCs, (GSE94820), ^47^), using CellHarmony ^48^ and SCmap ^49^ tools. Both mapping strategies (Figure 3a, Supplementary data Figure S3) confirmed, that the majority of LC single cell transcriptomes were distinct from the conventional DC1 (Figure 3a, 4%), and in contrast resembled cDC2 (Figure 3a, 64%) and cDC3 (Figure 3a, 15%). This is in agreement with the potent ability of both cDC2 and 3 to prime cytotoxic CD8 lymphocytes ^47^.

**Figure 3.**
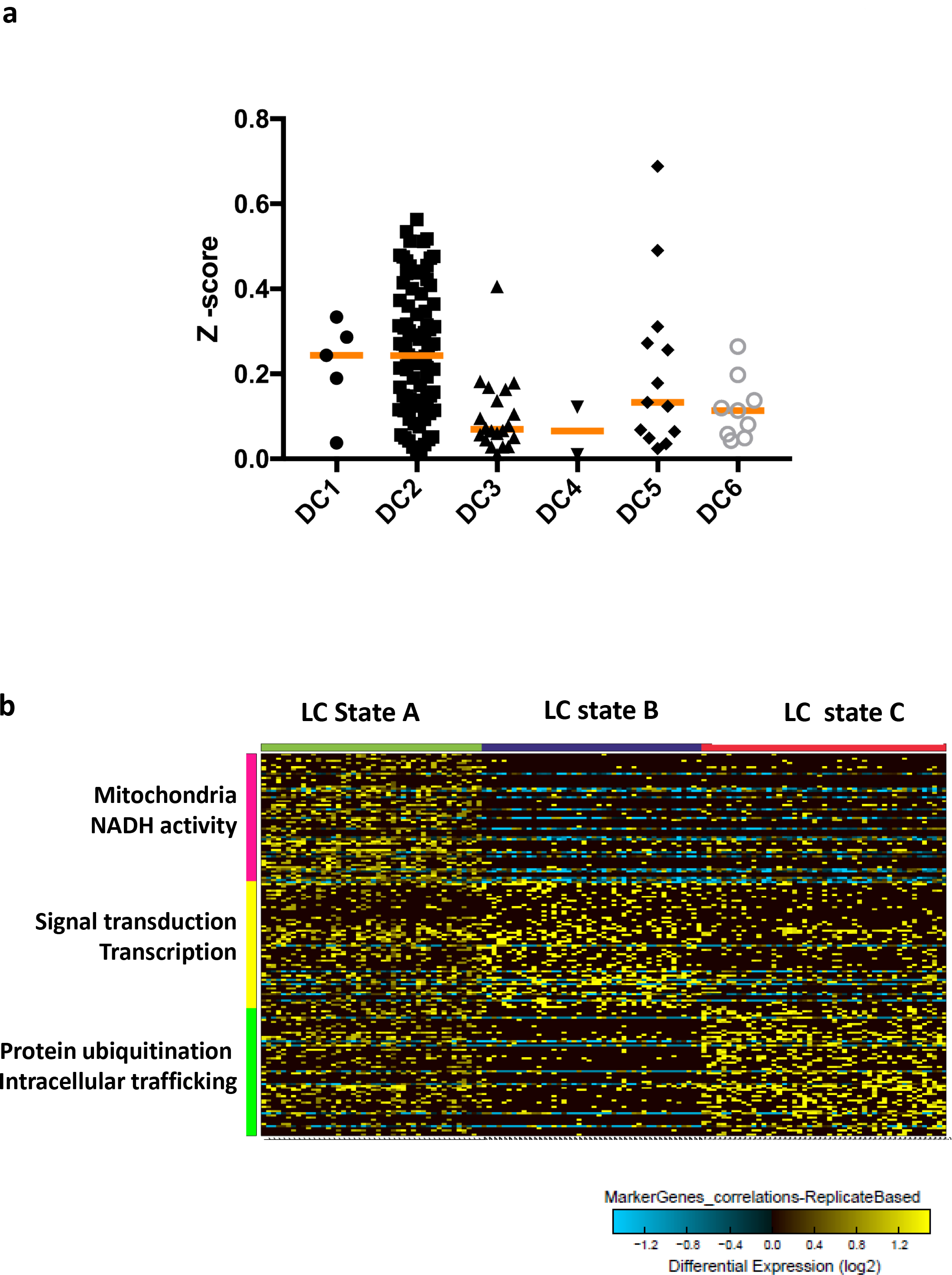
scRNA-seq maps LCs onto conventional DC2 and reveals three activation states. a. CellHarmony mapping of 138 single cell LC transcriptomes onto 6 blood DC sub-populations ^47^. Z Scores were calculated for each cell using AltAnalyze tool.
b. Euclidean correlation clustering (replicate based) of 181 marker genes in 138 single LCs. Analysis defined 3 distinct clusters varying in expression of genes involved in antigen processing and presentation (ICSG analysis, CPTT normalised counts, cosine clustering). Gene ontology analysis for marker genes was performed within AltAnalyzer. The heatmap shows log2 gene expression level.

Iterative clustering and guide-gene selection^50^ identified 3738 transcripts differentially expressed across the 138 LC cells (DEGs), grouped in three distinct clusters (Figure 3b). LC subset A was characterized by expression of genes involved in mitochondrial processes and NADH activity, subset B involved signal transduction and transcription while subset C included intracellular protein processing, ubiquitination and trafficking. Interestingly, while LC subset A showed increased expression of genes involved in all three of these categories, LC subset C, displayed expression of intracellular antigen processing genes with reduced levels of gene expression involved in mitochondrial metabolism and NADH activity. The coupling of oxidative phosphorylation with efficient priming of immune responses has been reported previously for murine DCs, increased catabolism was shown to be essential for DC tolerogenic function ^51^. Furthermore, mitochondrial membrane potential and ATP production enhances the phagocytic capacity of murine CD8 DCs, augmenting their cross-presentation at a late stage^52^. Thus, the LC subset A cells appear to be optimally programmed for priming of tolerogenic CD4 and protective CD8 T cell responses.

### LC migration and antigen cross-presentation is associated with induced expression of IRF4 and the absence of IRF8

Given critical roles for specific members of the IRF, ETS and AP-1 family transcription factors in antigen presentation in conventional DCs ^41,53^, we analysed the expression of IRF4, IRF8, PU.1, SPIB, cJUN, JUND and BATF3, in steady-state and migrated LCs. Whereas IRF4 and BATF3 proteins were expressed in steady-state LCs surprisingly there was no detectable expression of IRF8 (Figure 4a, Supplementary data Figure S4). Importantly, migration out of the skin further induced IRF4 expression (Figure 4a,b). In contrast, IRF8 protein remained undetectable in migrated LCs (Figure 4a). Analysis of transcripts for these transcription factors was in keeping with the expression of their proteins (Figure 4c, Supplementary data Figure S4a-e). We note that transcripts encoding IRF4, PU.1 (Spi1) and cJUN were most highly expressed in LCs and were at least an order of magnitude greater than those encoding IRF8 and the PU.1 paralog SpiB. Finally, TNF-α stimulation maintained the expression of IRF4 and BATF3. Importantly, IRF8 protein remained undetectable over the time course of TNF-α stimulation, in spite of low level up-regulation of IRF8 mRNA (Figure 4c, 4d, Supplementary data Figure S4f). Thus, antigen cross-presentation by human LCs appears to be IRF8 independent. This possibility was tested directly with murine LCs isolated from IRF8 knockout mice ^54^. IRF8-deficient LCs were able to cross-present the SIINFEKL epitope from the ovalbumin protein to OT1 cells, equivalently to their wildtype counterparts (Supplementary data Figure S4g-h). These results with murine IRF8 KO LCs are consistent with the findings with human LCs and demonstrate that antigen cross-presentation by LCs is indeed independent of IRF8. Instead, the results strongly suggest that human LCs depend on IRF4 along with PU.1 and BATF3 for programming expression of their cross-presentation genes.

**Figure 4.**
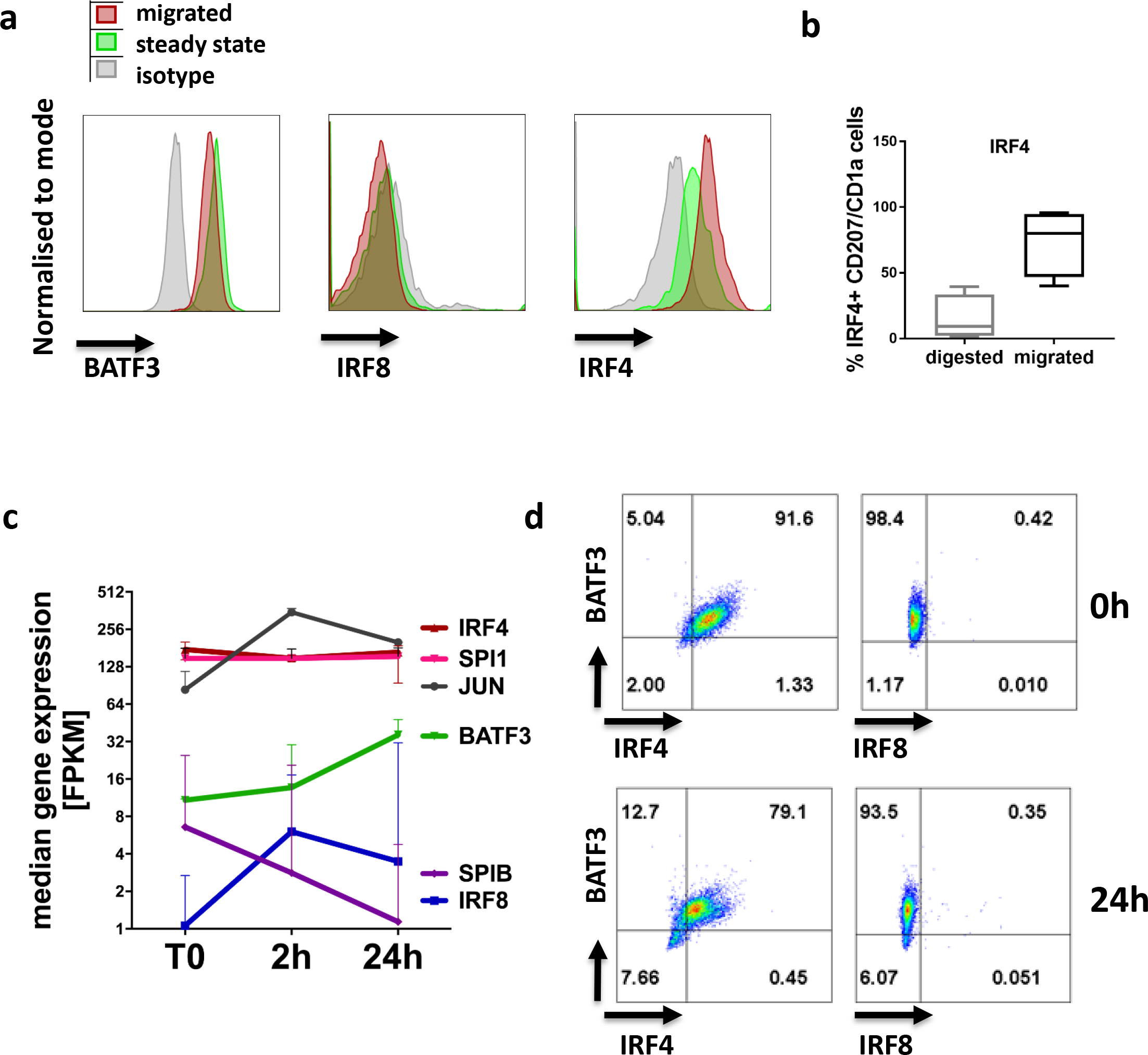
LCs induce expression of IRF4 upon their migration but lack IRF8. a. IRF4, but not IRF8 protein expression is induced in BATF3 positive LC during migration from the epidermis. A representative FACS analysis of 3-5 experiments, gates set using isotype controls for each antibody (Nuclear staining for IRF4, IRF8 and BATF3).
b. IRF4 protein expression in steady state vs migrated LCs. IRF4^+^ LCs (%) as measured by flow cytometry, median ± range, n=4, 3 paired samples.
c. Transcript levels of key transcription factors in steady-state migrated LCs before and after TNF-α stimulation (2h, 24h). FPKM values, median ± range of 3 biological replicates shown.
d. Human steady state migrated LC sustain expression of IRF4 and BATF3, after TNF-α signalling (24h). Representative graphs of 5 experiments, gates set using isotype controls for each antibody (Nuclear staining for IRF4, IRF8 and BATF3).

### IRF4 expression is correlated with the antigen cross-presentation programme in single cell LC transcriptomes

Having found that IRF4 is strongly induced when LCs migrate from the epidermis (Figure 4 a,b), a process associated with their enhanced antigen cross-presentation (Figure 1c), we wished to determine whether IRF4 likely regulates expression of various genes in the cross-presentation transcriptional module. Pairwise transcript correlation analysis was performed with IRF4 and all other DEGs (3737) in the LC sc-RNAseq dataset. This revealed 474 transcripts whose expression was positively correlated with IRF4 (r>0.1, Supplementary data Table S4). These included LC defining genes (*CD207*, *CD74*, *CD86*) and importantly, genes involved in antigen trafficking in the endosomal compartment as well as antigen processing, including ubiquitination (Figure 5a,b). In contrast, similar pairwise analysis of transcripts encoding IRF1, SPI1 and BATF3 with these IRF4 associated genes did not reveal a significant correlation (Figure 5c, Supplementary data 5a,b). Thus, IRF4 expression is uniquely correlated at the single cell level with the antigen processing, ubiquitination and proteasome degradation component of the antigen presentation and cross-presentation programme (Figure 5c, p<0.0001). Strikingly, ubiquitin-dependent protein catabolism was identified as the main gene ontology category enriched for 100% (89/89) of genes positively correlated with this signature (FDR p=9.9× 10E-134, Supplementary Table S3). It is therefore highly conceivable that IRF4, up-regulated during LC migration from the epidermis, specifically regulates expression of genes involved in protein ubiquitination in LCs, contributing to the ability of these cells to efficiently cross-present antigens.

**Figure 5.**
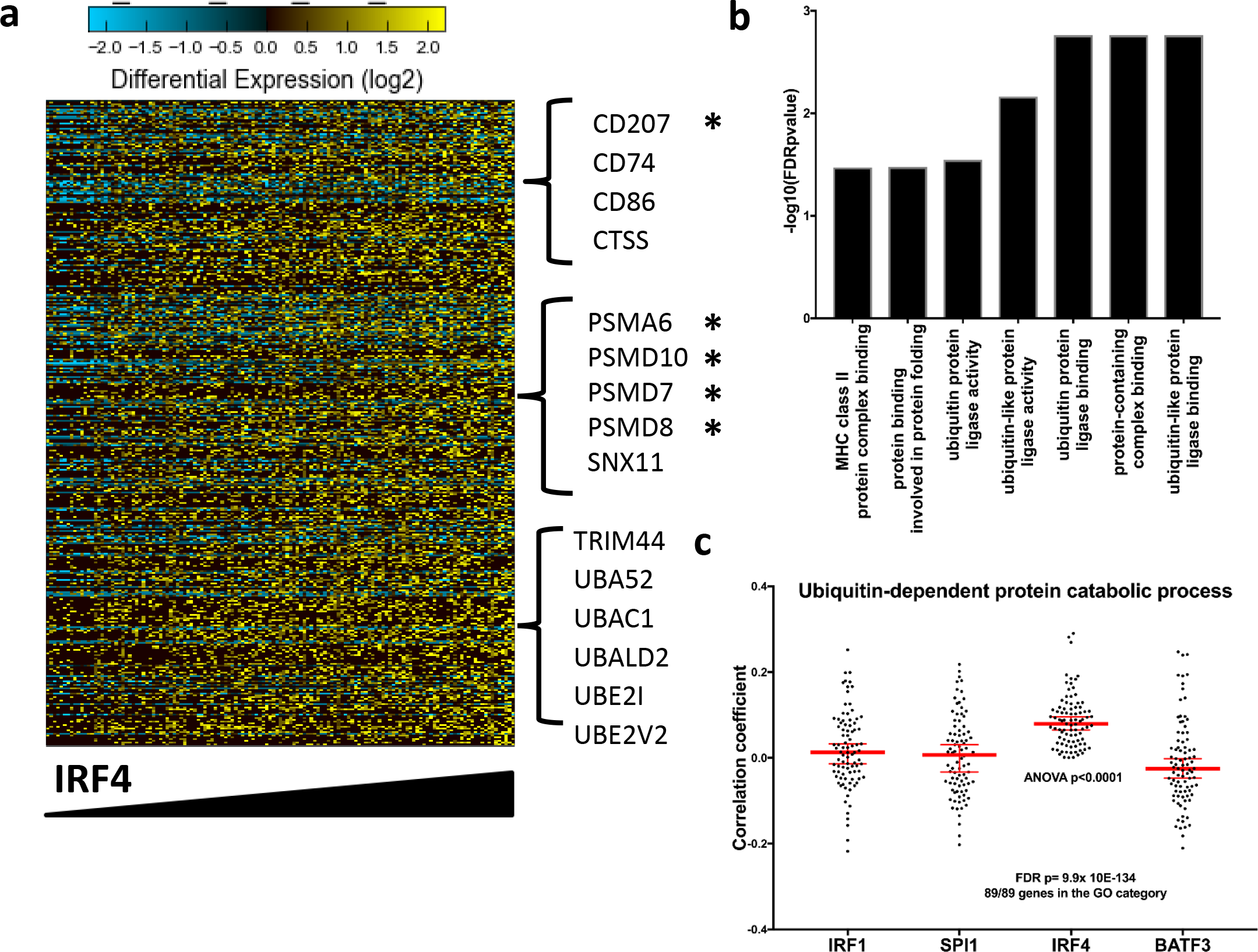
Single-cell iterative clustering analysis confirm dependence of LC antigen processing and presentation programme on IRF4 expression. a. Correlations between the CPTT values for DEGs and selected transcription factors were computed using corrplot and hmisc packages in R environment (Pearson coefficient >0.11,^92^). Cells are arranged according to increasing IRF4 transcript levels. Examples of genes correlated with IRF4 shown. Stars denote genes belonging to antigen presentation in class I, MSigBD, Broad Institute ^104^.
b. Gene processes enriched for IRF4 correlated genes were assessed using ToppGene. Top immune processes shown, significance denoted by FDR (Benjamini-Hochberg) corrected p-value.
c. Correlation coefficients across DEG involved in antigen processing and presentation in 138 single LCs were determined for indicated TFs. Pearson correlation coefficient is shown for each gene, median ± CI denoted in red. Gene ontology was assessed using ToppGene, significance denoted by FDR (Benjamini-Hochberg) corrected p-value.

### Enrichment of IRF composite elements in chromatin implicates IRF4-PU.1 and IRF4-BATF3 complexes in LC genomic programme for antigen cross-presentation

To uncover additional genes whose regulated expression could enhance the ability of migrated LCs for antigen cross-presentation, we performed RNA-seq in the absence of and with TNF-α stimulation for 2 or 24 hours. As shown above, TNF-α signalling leads to enhancement of LC antigen cross-presentation (Figure 1e). Although, the overall transcriptomic network remained relatively stable under these activation conditions, 1,156 genes were significantly differentially regulated by TNF-α (EdgeR, FDR<0.05, |LogFC|>1). Transcript-to-transcript clustering (BioLayout Express3D, r= 0.80; MCL = 1.7) identified 7 kinetic clusters with n>8 genes; 3 main clusters were characterised by gene expression peaks at 0, 2 and 24h (Figure 6a-c, Supplementary data Table S6). Gene ontology analysis indicated a consistent shift of the transcriptome towards a more activated LC phenotype; this included reduction of adhesion, enzymatic and trans-membrane signalling with the enhancement of immune functions (Supplementary data Table S6). Two waves of gene activation could be distinguished: an early wave, including the CD40 and CD83 genes, involved in T cell interactions and a late wave including PSME2 and TRIM21, 22 involved in antigen processing and protein ubiquitination (Figure 6a-c Supplementary data Table S6). In agreement with our microarray analysis ^22^, the late wave included components of immunoproteasome (*PSME2*, *ERAP2)*, genes involved in intracellular antigen trafficking between the cell membrane, the endosomal compartment and autophagosome (*CAV1*, *SQSTM1*) (Figure 6b). Interestingly, many members of the superfamily of tripartite motif-containing (TRIM) proteins were up-regulated in the late phase. TRIM proteins are E3 ubiquitin ligases involved in membrane trafficking, protein transport and protein degradation in proteasomes, crucial for many aspects of resistance to pathogens, and reported in the protection against lentiviruses ^55,56^. Thus, over the time course of stimulation with TNF-α, the LC transcriptome was highly enriched in genes involved in MHC I-dependent antigen presentation (Supplementary data Figure S6d).

**Figure 6:**
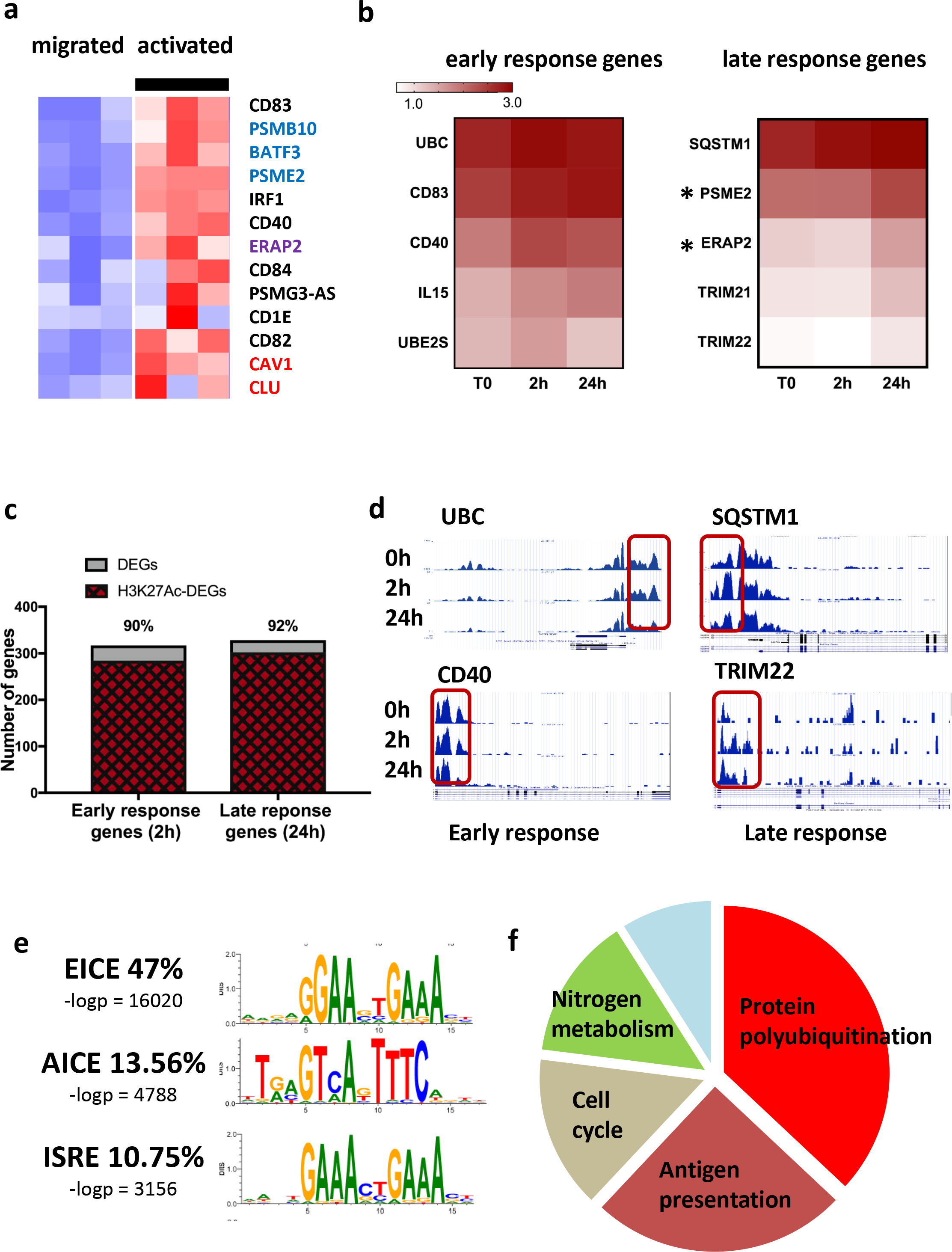
Chromatin landscape primes LC activation upon migration for antigen-presentation and is enriched for EICE, AICE and ISRE transcription factor motifs. Human steady state migrated LCs were subjected to whole genome chromatin profiling of H3K4me3 and H3K27Ac.

a. TNF-α stimulation (24h) of human LCs induces genes involved in antigen trafficking (red), processing (purple) and cross-presentation (blue). Median expression levels of 3 biological replicates (TMM normalised gene expression levels).
b. Enrichment of immune activation genes upregulated during a time course of TNF-α stimulation: left: early induced genes, peak expression at 2h, Cluster 3, right: late induced genes, peak expression at 24h. Median of 3 biological replicates, normalised expression levels. Stars denote genes belonging to antigen presentation in class I, MSigBD, Broad Institute ^104^
c. Proportion of DEG with H3K27Ac mark at 2h in clusters of co-expressed genes up-regulated early (2h, clusters 3) and late (24h, cluster 2) following stimulation with TNF-α. Changes in H3K27Ac acetylation were calculated using MANorm algorithm (MACS2, ^95^) embedded in BioWardrobe tool. Genes were filtered to include unique common entry across the biological replicates (n=3). Genes with detected changes in acetylation were intersected with DEGs identified by EdgeR analysis.
d. UCSC genome browser tracks of H3K27Ac mark changes in human migrated LCs over the timecourse of stimulation with TNF-α. Early ubiquitin C, UBC (top) and CD40 (bottom) and late sequestosome SQSTM1 (top) and TRIM22 (bottom) induced genes. Red rectangle denotes promoter site, a representative example.
e. Promoter sites of genes acetylated at H3K27 in steady-state human LCs are enriched in IRF-binding composite DNA elements. EICE is the top human motif enriched at steady state LCs (HOMER de novo motif detection analysis, median −logp value shown).
f. Peaks H3K4Me3 and H3K27Ac T0 datasets were scanned for ISRE/AICE/EICE binding motifs. 1193 consensus genes (present in all 3 biological replicates with both chromatin marks) were identified. Biological processes enriched in those genes were detected using ToppGene and collapsed using REVIGO (Similarity = 0.7, based on FDR corrected p-values for GO categories).

To explore if IRF4 along with its ETS and AP-1 family partners, PU.1 and BATF3 could be directly controlling the expression of antigen cross-presentation genes in human LCs, we performed genome-wide chromatin profiling and transcription factor motif enrichment analyses. This approach enables the inference of transcriptional regulatory sequences and the transcription factors that are acting to control gene activity in distinct cell types and cell states ^39,57^. Notably, tri-methylation at lysine 4 of histone 3 (H3K4Me3) marks active promoters ^58^, whereas acetylation of lysine 27 of histone 3 (H3K27Ac) marks active transcriptional enhancers; the latter have been postulated to be the primary determinants of tissue-specific gene expression ^59,60^. Thus, we performed ChIP-seq to analyse the genome-wide landscape of H3K4Me3 and H3K27Ac in migrated and TNF-α-activated LCs. Three independent samples of human migrated LCs exhibited a highly conserved histone modification landscape across the genome with 95% of H3K4Me3 marks in the same regions (Supplementary Table S4). Analysis of H3K27 acetylation peaks in the same LC preparations showed 78% to be shared across all three samples (Supplementary data Table S5). Of 13,402 H3K4Me3 peaks common for steady state LCs (11,665 unique genes), over 92% of the H3K4Me3 peaks were mapped to promoter regions (Supplementary data Table S4). In contrast, while 62% of H3K27Ac peaks were positioned either within the promoter region or upstream, a significant proportion of these peaks were distributed in other areas of the genome (Supplementary data Table S5). This distribution was expected for intergenic or intronic enhancers that function at large distances from the promoters on which they act.

Focusing the analysis on genes associated with immune function (InnateDB: Immunome collection,^61^), we identified 290 immune genes with active (H3K4Me3) promoters. These were highly enriched in genes encoding receptor binding and activation (in particular able to respond to TNF-α cytokine family signalling (FDR p= 1.2×10E-12), and genes involved in antigen processing and presentation (FDR p = 10E-22). As noted above, considerable overlap between H3K4Me3 and H3K27Ac marks was apparent at promoter regions. This indicated that the migrated LCs are pre-programmed for efficient antigen processing and presentation (Figure 6 b-d). In concordance with the observed gene expression pattern, histone marks were very low or absent on genes involved in innate inflammatory responses, such as production of cytokines (Supplementary data, Table S4, S5). This was in marked contrast to genes involved in antigen presentation. To analyse whether genomic programming of human steady-state LCs was similar to other known cell types including monocytes, DCs and macrophages, we compared our ChiP-seq profiles to a large collection of publically available genomics datasets (see Methods). Surprisingly, the chromatin landscape of steady-state migrated LCs was strikingly similar to that of macrophages and CD14+ monocytes, and significantly less strongly correlated with that of dendritic cells (Supplementary Table S6).

TNF-α-induced alterations in the chromatin landscape of LCs correlated with corresponding directional changes in TNF-α-regulated genes. Thus, stimulation with TNF-α induced up-regulation of H3K27Ac acetylation in genes underpinning LC activation (Figure 6c,d). Two hours after TNF-α exposure, the high levels of H3K27 acetylation, observed in steady state LCs (Figure 6c) were consistently maintained for 90% and 92% of genes induced during the early (cluster 3) and late (cluster 2) waves of responses, respectively (Figure 6c). Moreover, TNF-α signalling enhanced H3K27Ac levels in over 50% of genes (Supplementary data Figure S6 e,g). In line with the transcriptome changes, genes with enhanced H3K27Ac marks at 2h encoded innate immune processes including leukocyte activation (FDR p-value = 2×10E-19) and co-stimulation (FDR p-value = 7.99×10E-16). A significant proportion of genes (286) within this programme was associated with ubiquitin-mediated proteolysis, highlighting the importance of this process for LC function (FDR p = 3.8 × 10E-85). In contrast, genes involved in cell cycle and motility were characterised by reduced histone acetylation marks. Consistent with their unstimulated profile, and the lack/low levels of mRNA, many pro-inflammatory cytokine loci (such as IL1b, IL12a), classically associated with DC immune activation, were characterised by paucity of histone marks upon stimulation with TNF-α, both with regard to H3K4Me3 and H3K27Ac. It is worth noting that the majority of histone marks induced by TNF-α signalling were readily detected prior to activation (Figure 6c,d), suggesting that the migrated LCs were pre-programmed for rapid immune activation from a genomic standpoint.

To test if IRF4 along with its ETS and AP-1 family partners, PU.1 and BATF3 could be directly controlling the expression of antigen cross-presentation genes in human LCs, we performed transcription factor (TF) motif enrichment analyses. There was extremely high enrichment of the known composite IRF-binding sequences ^35^ at the promoters of genes carrying H3K4Me3 and H3K27Ac marks (Figure 6e). The ETS-Interferon Composite Element (EICE) was the most frequent and significantly enriched TF motif in steady state migrated LCs (47%, −logp=16,020), while the AP1-Interferon Composite Element (AICE) and the interferon-stimulated response element (ISRE/IRF1) were present at frequencies of 13.6% (-logp=4788) and 10.3% (-logp=3.156), respectively (Figure 6e, Supplementary Table S8). Analysis of biological processes enriched for genes carrying H3K4Me3 and H3K27Ac marks in their promoter regions and predicted IRF4 binding motifs demonstrated, that over 60% were involved in protein polyubiquitination, or antigen processing and presentation (Figure 6f). Thus, transcription and chromatin profiling coupled with TF motif analysis strong suggests that IRF4 in conjunction with PU.1 and BATF3, binding to EICE and AICE motifs, respectively, program the expression of a large set genes in human LCs that are involved in antigen cross-presentation.

## Discussion

Since their discovery by Paul Langerhans in 1864, LCs have been puzzling scientists. Despite being the longest studied antigen presenting cell population, and being considered the stereotypical “dendritic cells”, their place in the innate immune cell spectrum and developmental programming remain elusive. This conundrum is in part caused by the difficulties in studying LC, arising from paucity of *in vitro* models, controversy around defining a suitable LC progenitor, and the differences between human and murine skin immunology ^62^. To address these challenges in the analysis of primary human LCs, we utilized an integrative systems immunology approach, combining transcriptome analysis at the population and single cell levels with profiling of the chromatin landscapes ultimately coupling these genomic profiles with LC phenotypic and functional characteristics. While our investigations are restricted by the availability of the cells and the limitations of experimental models, we believe that our approach offers the opportunity of addressing some of the key molecular questions in primary human LCs, enabling us to propose a plausible model for regulation of LC activation. Our extensive transcriptomic and epigenomic analyses of primary human skin-derived LCs reveals three striking features: (i) The pre-programming and relative stability of the LC chromatin landscape and transcriptome (ii) the importance of migration from the epidermis for activation of the LC transcriptome programme and (iii) the likely dependence of LC migration and function, in particular their ability to cross-present antigens, on a major immune transcriptional regulator IRF4 rather than its paralog IRF8. All three features are likely to be interconnected, and at least in part dictated by the localisation of LCs in the epidermis ^63^.

By profiling the H3K4Me3 and H3K27Ac histone modifications across the LC genome, we were able to construct the chromatin landscape of human migrated LCs for the first time, and document that their transcriptomic programme encoding antigen presentation genes is poised for efficient expression during, or before their migration from the epidermis. TNF-α, a pro-inflammatory signal produced in the skin plays a critical role in activating LC emigration from the epidermis, hence, in inducing LC immunogenic function and ability to present antigens. Stimulation with TNF-α, significantly enhanced the pre-existing transcriptional programme, further confirming that the LCs are fully committed for efficient antigen presentation and cross-presentation. Such genomic programming, realised at the level of transcriptional enhancers, could be both developmentally and environmentally specified. The latter, termed “trained immunity”, has been recently documented in murine myeloid cell progenitors during vaccination with BCG ^64^, where the vaccine induced persistent H3K27Ac marks in regions in macrophages that enriched for genes responding to a subsequent *M. tuberculosis* infection. We propose that the chromatin landscape observed in adult LCs is initiated from the earliest stages of their embryonic development and then overlaid by exposure of LC precursors and LCs to signals from the tissue microenvironment as well as environmental exposures, the latter from symbiotic microbial colonisation, vaccines and infections.

We have delineated three distinctive IRF-binding motifs, EICE, AICE and ISRE as key regulatory elements associated with expression of the LC transcriptomic programme. Classically, EICE and AICE have been shown to be bound by IRF4 or IRF8, in combination with their transcriptional partners from either ETS or AP-1 transcription factor families ^35,65–68^. Our analysis of IRF4 and IRF8 protein expression and their transcriptional binding partners clearly demonstrates that IRF8 protein is not detectably expressed in human LCs and is thus dispensable for their ability to cross-present. This conclusion was strengthened by the demonstration that murine LCs lacking IRF8 cross-present antigens as efficiently as their wild type counterparts. The high levels of IRF4 protein which is induced in LCs upon migration make it a likely candidate as a transcriptional regulator of cross-presentation. In cDC1, one of two main blood DC subsets, cross-presentation has been shown to depend critically on IRF8 and BATF interaction with composite elements on the promoters of key target genes. By contrast, recent reports in other cell types, such as MoDCs ^69^, cDC2 (Murphy, personal communication) and red pulp macrophages ^70^ indicate that the same transcriptomic programme can be successfully initiated and executed by IRF4 ^69^. Hence the high levels of IRF4 expressed in human migrated LCs, are likely to be involved in the orchestration of this programme ^69^, enabled by efficient DNA binding to either EICE or AICE composite elements in combination with PU.1 or BATF3, respectively. Both of these transcription factors are co-expressed in LCs with IRF4. We note that IRF4 can also bind directly to ISREs as a homodimer at high concentrations ^68^ and that this motif was also statistically enriched in the LC chromatin profile.

The argument for a key role for IRF4 in orchestrating the ability of LCs to cross-present is further strengthened by our analysis of gene expression at the single cell level. While expression of all the transcription factors examined is correlated with the cross-presentation programme, IRF4 is exclusively and highly significantly correlated with expression of genes involved in protein ubiquitinase activity; this gene module is distinguished in LCs upon their activation through chromatin modifications and at the transcriptome level. Interactions between antigen presentation and protein ubiquitination have been extensively investigated, and several modes of interaction can explain the importance of the latter process for antigen cross-presentation. Ubiquitin ligases such as MARCH9 and Sec61 have been directly implicated in facilitating cross-presentation ^71,72^, while other members of the MARCH family enhance phagocytosis and antigen presentation through ubiquitination of MHC molecules ^70^ or facilitate entry of MHC molecules to the endoplasmic reticulum (ER) ^73^. Since abrogation of polyubiquitination strongly reduces the epitope presentation from ER-processed antigens ^74^, it is conceivable that increase of polyubiquitination expands the spectrum of epitopes in activated. The notable lack of IRF8 expression in LCs, consistent with inactive chromatin at the IRF8 locus, distinguishes LCs from both macrophages and DCs and likely contributes to their discrete role in epidermal homeostasis. Independence of LCs from IRF8 could represent a mechanism for their plasticity, enabling them to be adapted to their environmental niche. IRF8 has been shown to regulate production of pro-inflammatory cytokines in DCs ^75,76^, and macrophages, the latter contributing to chronic inflammation ^77^. We propose that by utilising IRF4 rather than IRF8, LCs uncouple the cross-presentation of antigens from production of pro-inflammatory mediators, and thus prevent excessive inflammatory responses and promote epidermal homeostasis.

The epidermis forms a unique environmental niche, constantly exposed to insults from the outside world and to interactions with symbiotic and pathogenic microbiota. Inhabiting this niche, LC need to react rapidly to any changes and hence require highly efficient transcriptomic networks in order to initiate precise and efficient adaptive immune responses. In contrast, excessive pro-inflammatory signals need to be limited, in order to preserve tissue homeostasis. At the same time, due to their external localisation, LCs are constantly exposed to environmental insults, and thus need to develop resilience to DNA damage and re-structuring, as described in the “access-repair-restore” model ^78^. Relative stability of LC transcriptomic networks and chromatin landscape very efficiently meets the criteria required, preventing both excessive inflammatory signalling and accumulation of molecular alterations due to the interactions with environment. Cross-talk with keratinocytes has been shown to be fundamental for LC development and keratinocyte-derived TGF-β and BMP7 have long been recognised as critical for LC development both in vivo and in vitro ^15,79,80^.

Recent studies investigating the interactions between LCs and keratinocytes highlight the importance of E-cadherin and alpha-v-beta-6 integrin signalling for LCs function reflected in immunisation responses to an epicutaneous protein antigen ^81,82^. It is conceivable, that this intimate cross-talk prevents LC activation *in situ*, preserving epidermal homeostasis. On migration, such cross-talk between LC and keratinocytes is broken, resulting in the initiation of LC immune programmes. Two elements act in concert in this process: programming of the LC chromatin landscape and transcription factor expression. Our results show that indeed, from the very earliest time point following migration from the skin, antigen presentation by LCs is poised by chromatin modification. The same migration efficiently stimulates expression of IRF4, and co-expressed transcriptomic programmes encoding antigen uptake, processing and presentation. It is highly plausible that the increase in IRF4 expression during LC migration out of the epidermis makes LCs immune-stimulatory, activating IRF4-coordinated transcriptomic programmes centred around efficient antigen presentation and ubiquitination. Synergistically with LC migration, pro-inflammatory cytokine signalling enhances LC ability to process antigens. Interestingly, genes encoding protein ubiquitination and ubiquitin-dependent proteolysis were amongst the most responsive to TNF-α. While our analyses point to coordinated expression of IRF4 and genes encoding ubiquitin-dependent proteolysis, IRF4 protein levels remain stable during stimulation with TNF-α. In contrast, levels of H3K27Ac around the promoter regions of those genes increases, potentially enhancing their transcription. It is therefore conceivable that coordinated action of IRF4 and chromatin landscape modification determines the ability of LC to rapidly increase antigen presentation and cross-presentation in the event of infection or inflammation. These observations lead us to propose a two-signal model of LC activation, where loss of contact with epidermal structural cells initiates LC activation programme and induces IRF4 expression. Subsequent (or concurrent in an infection setting) pro-inflammatory cytokine signalling from other cutaneous cell types stabilises the LC transcriptomic programme and enhances efficient activation of adaptive immune responses.

## Methods

### Cell isolation and stimulation with TNF-α

#### Human LCs

Skin specimens and blood samples were acquired from healthy individuals after obtaining informed written consent with approval by the Southampton and South West Hampshire Research Ethics Committee in adherence to Helsinki Guidelines [ethical approvals: 07/Q1704/59, NRES 07 Q1704 46]. Split skin was obtained using graft knife and subjected to dispase (2U/ml, Gibco, UK, 20h, +4℃) digestion of epidermal sheets. Migrated LCs were harvested after 48 hours culture of epidermal fragments in full culture media (RPMI, Gibco, UK, 5%FBS, Invitrogen, UK, 100 IU/ml penicillin and 100 mg/ml streptomycin, Sigma, UK). Low density cells were enriched using density gradient centrifugation (Optiprep 1:4.2, Axis Shield, Norway ^83^ and purified with CD1a+ magnetic beads according to manufacturer’s protocol (Milenyi Biotec, UK). Steady-state migrated LCs were processed for RNA-seq and ChIP-seq experiment or immediately cryopreserved in 90% FBS (Gibco, UK), 10% DMSO (Sigma, UK). For genomic and transcriptomic analyses of activated LCs fresh migrated LCs from 3 donors were stimulated with TNF-α (25 ng/ml, Miltenyi Biotec, UK) for 2, and 24 hours (RNA-seq: 3 × 10^5^ cells/donor/time point, ChIP-seq: 1.5-2 × 10^6^ cells/donor/time point, paired samples from the same donor for RNA-seq and ChIP-seq). Steady-state LCs were enzymatically digested from the epidermal sheets using Liberase™ TM research grade (Roche, UK, 2h at 37℃).

#### Murine LCs

Wild type (wt) and IRF8-/- LCs were isolated from ears and tails from 3 mice/type maintained on C57Bl/6 background. Mice were bred and housed by Cincinnati Children’s Hospital Medical Center (CCHMC) Veterinary Services, and mouse manipulations were reviewed and approved by the Children’s Hospital Research Foundation Institutional Animal Care and Use Committee (Protocol Number IACUC2013-0090). Dorsal and ventral sides were split with tweezers, and the epidermis was isolated following dispase (2U/ml, Gibco, UK, 20h, +4℃) digestion. Pooled wt and IRF-/- epidermal migrated cells were enriched on Lymphoprep gradient (Diagenode, USA), and used for the flow cytometry staining and cross-presentation experiments.

### Antigen cross-presentation assay

#### Human LCs

Cells were pulsed with 10 *μ*M proGLC (FNNFTVSFWLRVPKVSASHLEGLCTLVAML; Peptide Protein Research, Fareham, UK) for 24 hours, supplemented with TNF-α (25 ng/ml, Miltenyi Biotec, UK) after initial 2 hours. Human responder cells: PBMC from HLA-A2 individuals were isolated by Ficoll-Hypaque density gradient centrifugation and co-cultured with 40 μM EBV peptide for 12 days in complete medium supplemented with 1% sodium pyruvate (Gibco, UK) plus 10% human serum (Sigma,UK). IL-2 (100 IU/ml, Peprotech, UK) was added every 3 days. IL-2 was removed from the culture for 24 hours prior to testing in ELISpot. For ELISpot assays, TNF-α matured and washed EBV peptide pulsed LCs (1×10^3^ cells) were co-cultured with GLC peptide-specific T cells (5×10^4^ cells/ per well) for 20 hours as per manufacturer’s protocol (Mabtech, Sweden).

#### Murine LCs

Enriched migrated LCs were pulsed with ova protein (100 *μ*g/ml, Pierce Chemical Co, USA) for 24h, supplemented with TNF-α (50 U/ml, Peprotech, USA) after initial 2 hours. Murine responder cells: OT I cells were freshly prepared from OT I mice spleen by mechanical digestion, red cell removal (ACK lysing buffer, Quality Biological, USA), followed by naive CD8 T cell purification by magnetic-activated cell sorting accordingly to the manufacturer protocol (MACS, Miltenyi Biotech, USA). *Ova*-pulsed and control, wt and *IRF8-/-* LCs were incubated with OT I cells (20,000 LCs: 10^6^ T cells) for 48h, supplemented with brefeldin A (BD Biosciences, USA) for the last 6h before testing with flow cytometry for intracellular IFN-γ production. Cells were permeabilised with Cytofix/Cytoperm kit (BD Biosciences, USA) accordingly to the manufacturer protocol and stained with monoclonal antibodies targeting IFN-γ (BD Bioscience, USA). Analysis was performed on live AQUA-negative (Invitrogen, USA), CD3/ CD8 (BD Biosciences, USA) expressing cells.

Flow cytometry: All antibodies were used at pre-titrated, optimal concentrations. For surface staining of live cells buffer containing 5% FBS and 1% BSA was used for all antibody staining. FACS Aria flow cytometer (Becton Dickinson, USA) was used for analysis of human LCs for the expression of CD207, CD1a, HLA-DR (mouse monoclonal antibodies, CD1a, CD207:Miltenyi Biotech, UK and HLA-DR: BD Biosciences, UK). For transcription factor intra-nuclear staining cells were permeabilised with Foxp3 / Transcription Factor Staining Buffer Set (eBiosciences, UK) accordingly to the manufacturer protocol, and stained with monoclonal antibodies targeting IRF4, IRF8, BATF3, PU.1, cJUN, (IRF4:rat monoclonal, eBiosciences, UK, mouse monoclonal: IRF8, eBiosciences, BATF, R&D Systems, JUN Millimark, UK, PU.1 Biolegend, UK). IRF1 staining was done using rabbit polyclonal anti-human IRF1 antibody (Abcam, UK) following fixation with 80% methyl-alcohol and permeabilisation with Tween20. Analysis was performed on live AQUA-negative (Invitrogen, UK), CD207+/HLADR+ migrated or steady-state LCs, in comparison with appropriate isotype controls.

### RNA-seq

RNA was isolated using RNeasy mini kit (Qiagen) as per the manufacturer’s protocol. RNA concentration and integrity was determined with an Agilent Bioanalyser (Agilent Technologies, Santa Clara, CA). All the samples had a RNA integrity number of 7.0 or above and were taken forward for labelling. RNA-seq libraries were generated from 300 ng total RNA with an RNA Sample Prep Kit (Illumina) according to a standard protocol. The libraries were sequenced with Illumina HiSeq2500 in the DNA sequencing core of the Cincinnati Children’s Hospital Medical Center. Each sample was used to generate 2 × 10^7^ reads with 75– base pair paired-end sequencing.

### ChIP-seq

Purified migrated LCs were fixed with 1% formaldehyde for 15 min and the reaction quenched with 0.125 M glycine. Chromatin was isolated by the addition of lysis buffer, followed by disruption with a Dounce homogenizer. Lysates were sonicated and the DNA sheared to an average length of 100–200 bp (Covaris). Genomic DNA (Input) was prepared by treating aliquots of chromatin with RNase, proteinase K and heat to remove crosslinks, followed by ethanol precipitation. Pellets were resuspended and the resulting DNA was quantified on a NanoDrop spectrophotometer. Extrapolation to the original chromatin volume allowed quantitation of the total chromatin yield. Genomic DNA regions of interest were isolated using 2.8 μg of antibody against H3K27Ac or H3K4Me3 ^84^. Complexes were washed, eluted from the beads with SDS buffer, and subjected to RNase and proteinase K treatment. Crosslinks were reversed by incubation overnight at 65 °C, and ChIP DNA was purified by phenol–chloroform extraction and ethanol precipitation. Illumina sequencing libraries were prepared from the ChIP and input DNAs by the standard consecutive enzymatic steps of end-polishing, dA-addition, and adaptor ligation using Truplex™ -FD prep kit (Rubicon Genomics, USA). After a final PCR amplification step, the resulting DNA libraries were quantified using Qubid and 75 nucleotide single-end reads were sequenced Illumina HiSeq2500 in the DNA sequencing core of the Cincinnati Children’s Hospital Medical Center.

### Drop-seq

Highly purified human LCs (> 80% of CD1a+HLADR+) were unbanked from cryo-storage, and processed on ice to enable the co-encapsulation of single cells with genetically-encoded beads (Drop-seq, ^85^). Monodisperse droplets at 1 nl in size were generated using the microfluidic devices fabricated in the Centre for Hybrid Biodevices, University of Southampton. To achieve single cell/single bead encapsulation barcoded Bead SeqB [Chemgenes, USA], microfluidics parameters (pump flow speeds for cells and bead inlets, cell buoyancy) were adjusted to optimise cell-bead encapsulation and the generation of high quality cDNA libraries. Following encapsulation, ~4500 STAMPS (beads exposed to a single cell) from 1.2 ml cell suspension were generated in the Faculty of Medicine University of Southampton Drop-seq Facility. An estimated 150 STAMPS (based on encapsulation frequencies and bead counts) were taken further for library prep (High Sensitivity DNA Assay, Agilent Bioanalyser, 12 peaks with the average fragment size 500 bp). The resulting libraries were run on a shared NextSeq run (1.5×10E5 reads for maximal coverage) at the Wessex Investigational Sciences Hub laboratory, University of Southampton, to obtain single cell sequencing data.

### Bulk RNA-seq data analysis

Quality control for FASTQ files with raw sequence data was done using FASTQC tool [FastQC: a quality control tool for high throughput sequence data. Available online at: http://www.bioinformatics.babraham.ac.uk/projects/fastqc] Adapter sequences and low quality reads were trimmed using Trimmomatic ^86^. High-quality reads were mapped to the human genome (hg19) using TopHat (version 2.0.9, ^87^) and, following the removal of multimapping reads, converted to gene specific read counts for annotated genes using HTSeq-count (version 0.5.4) ^88^. Raw counts from RNA-Seq were processed in Bioconductor package EdgeR ^89^, variance was estimated and size factor normalized using TMM. Genes with minimum 2 reads at minimum 50% samples were included in the downstream analyses. Differentially expressed genes (DEG) we identified applying significance threshold FDR p<0.05, |LogFC|>1. Normalised reads from DEG were taken for transcript-to-transcript co-expression analysis (BioLayout *Express*^3D^, ^90^. FPKMs were estimated using Cufflinks package ^87^. Gene ontology analysis was done using ToppGene on-line tool ^91^.

### scRNA-seq data analysis

Digital gene expression data was generated for the 150 STAMPs following the pipeline established in the McCarrol lab ^85^. Samples were de-multiplexed with bcl2fastq tool from Illumina, and tag bam files with read sequence extended were processed to differentiate molecular barcodes, cell barcodes and genome reads. Reads with low quality bar codes were removed, and the unpaired reads were aligned with STAR. To create a matrix of digital gene expression data, the total number of unique UMI sequences was counted for each transcript for a given cell ^85^. Highly expressed genes and high-quality cells were selected through filtering by both the mean count of each gene across all cells and also the mean count of all genes expressed in a single cell (mixture model filtering). This reduced the matrix from 15200 genes and 150 cells to 9075 genes and 118 cells. To investigate cell-to-cell variability in gene expression Hierarchical clustering (clust=ward.D2, dist = canberra) was applied. Transcript-to-transcript correlation was assessed using BioLayout *Express*^3D 90^, corrplot and hmisc packages in R environment ^92^. Differentially expressed genes were identified through comparing mean gene expression values between clusters of interest. Genes with a |logFC| >1 in expression between comparisons were submitted to gene ontology analysis in Toppgene (adj. p-value <0.05) to characterise the specific biological processes and pathways associated with each subpopulation. To assess gene expression signatures and pathway activation, a non-parametric and unsupervised software algorithm called GSVA software in the R package with the RNA-seq mode was used ^93^. We included a set of marker genes from monocytes and DC from Villani et al. as a reference set ^47^. Over-representation analysis was performed using the hypergeometric test. The results were plotted as a heat map using hierarchical clustering.

### ChIP-seq data analysis

ChIP-seq data analysis was performed using pipelines implemented in the BioWardrobe suite ^94^ BAM-formatted hg19 aligned ChIPseq reads were used for peak calling MACS2 (version 2.1.1. ^95^ was used to estimate fragment size, identify and annotate peaks. Changes in histone modification profiles were assessed with MAnorm algorithm implemented in BioWardrobe. ChiP-seq profiles at a distance −3000bp - +3000bp from TSS were generated using ChIPseeker package in R. Common regions with histone modification were identified using findoverlaps function, DiffBind package (R environment.

### Transcription factor binding site motif enrichment analysis

We performed TF motif enrichment analysis on ChIP-seq peak sets using the HOMER sortware package ^96^. HOMER calculates the statistical enrichment in a set of input DNA sequences (here, ChIP-seq peak sequences) for a large set of position weight matrix (PWM) TF binding models. Subsequently, putative binding sites for enriched motifs (EICE, AICE, and ISRE) were identified in the H3K27ac T0 dataset using custom scripts encoding the standard log likelihood PWM scoring function ^97^. DNA sequences scoring at least 70% of the best possible log-likelihood score were recorded as putative binding sites.

### Intersection of ChIP-seq peaks with available public functional genomics datasets

To identify transcription factor binding events and other epigenetic marks that overlap with our ChIP-seq data, we used our Regulatory Element Locus Intersector (RELI) computational method ^98^, which overlaps a set of genomic locations (e.g., peaks from a ChIP-seq experiment) with a large collection of functional genomics datasets. We created a library of ~5,000 datasets by compiling data (ChIP-seq for TFs and histone marks, DNase-seq, ATAC-seq, etc.) from a variety of sources, including ENCODE ^99^, Cistrome ^100^, PAZAR ^101^, Re-Map ^102^, and Roadmap Epigenomics ^103^. As input, RELI takes a set of genomic loci (e.g., H3K27ac ChIP-seq peaks at T0). These loci are systematically intersected with each functional genomics dataset, and the number of input regions overlapping each dataset by at least one base are counted. Next, a p-value describing the significance of this overlap is estimated using a simulation-based procedure. To this end, a ‘negative set’ is created for comparison to the input set, which is created by compiling all regions of open chromatin in the genome (i.e., the union of all available human DNAse-seq, ATAC-seq, and FAIRE-seq datasets). A distribution of expected overlap values is then created from 2,000 iterations of randomly sampling from the negative set, each time choosing a set of negative examples that match the input set in terms of the total number of genomic loci and the length of each locus. The distribution of the expected overlap values is used to generate a Z-score and corresponding p-value estimating the significance of the observed number of input regions that overlap each data set. Collectively, this procedure controls for the count and sizes of the input loci, and the count and sizes of each individual dataset in the library.

#### Accession codes

Sequencing data for RNA-seq, scRNA-seq and ChiP-seq is stored in Gene Expression Omnibus database, submission number GSE120386.

## Extended data

**Figure S1. Phenotype and activation status of primary human LC**

a) Flow cytometry assessment of steady-state and migrated LCs. LCs were enumerated as CD207/CD1a/HLA-DR high cells. Expression of co-stimulatory molecules critical for CD8 T cell activation during antigen presentation was measured. A representative example of n>5 experiments.

f) IFN-γ secretion by EBV-specific CD8 T cell line stimulated with antigen presenting LCs in the context of MHC I HLA-A2. Steady-state or migrated LCs were pulsed with 9 amino acid peptide GLC, an EBV epitope (GLC, dark grey), or unpulsed (light grey). Pulsed or unpulsed (light gray) LCs were stimulated with TNF-α and then assayed for IFN-γ secretion. ELISpot assay, n=5 independent experiments in triplicate.

**Figure S2 Comparative analysis of steady-state human LCs and other cross-presenting DC subsets**

a. Overlaps between reported cross-presentation signatures: Reactome database (76 genes), Artyomov et al (311 genes) and genes expressed in migrated LC >10 FPKM.
b. Heatmap of log(2) FPKM gene expression levels for genes involved in antigen processing and presentation in class I, MSigDB, Broad Institute ^104,105^

**Figure S3. Analysis of single cell RNA-seq in human migrated LC**

150 single migrated epidermal cells highly enriched in LC were subjected to Drop-seq encapsulation and single cell RNA-sequencing. Reads were aligned against human reference genome (hg19) using STAR aligner (v2.5.0a), then sorted/converted/merged to a BAM with a tag “GE” onto reads for data extraction. The DigitalExpression program (Dropseq-tools v1.0) performed digital counting (DGE) of the mRNA transcripts (unique molecular identifiers to avoid double counting reads/PCR duplicates) and created a DGE matrix (one measurement per gene per cell). Cells expressing less than 300 or more than 3,000 genes were excluded from further analysis. Digital gene expression matrix was subjected to Seurat analytical pipeline, to identify cell subpopulations and cluster defining markers.

a. Purity of migrated epidermal cells subjected to single cell analysis, 83% cells were CD1a/HLA-DR positive, expressing classical migrated LCs phenotype,
b. tSNE plot identifying contamination of epidermal LCs (red) with melanocytes (blue)
c. Expression of selected cluster defining markers identified with Seurat analysis across cell population.
d. scRNA_seq LC data was normalized using SCnorm ^106^ in R version 3.5.1; Projection onto a reference set of newly classified dendritic cells ^47^ was carried out using Gene Set Variation Analysis (GSVA) ^93^ based on enrichment score. The heatmap shows the GSVA enrichment score for each LC transcriptome.
e. The heatmap shows the Pearson correlation coefficient for each LC transcriptome.

**Figure S4. Transcription factor expression in steady-state and migrated LCs**

a. Flow cytometry analysis of intranuclear IRF4 protein expression in migrated and steady-state LCs (CD1a/CD207^high^). Gates set using isotype controls. Data representative of 8 independent skin donors
b. Flow cytometry analysis of intranuclear IRF8 protein expression in migrated and steady-state LCs (CD1a/CD207^high^). Gates set using isotype controls. Data representative of 3 independent skin donors
c. Flow cytometry analysis of intranuclear PU.1 protein expression in migrated and steady-state LCs (CD1a/CD207^high^). Gates set using isotype controls. Data show an independent experiment (n=1)
d. Flow cytometry analysis of intranuclear BATF3 protein expression in migrated and steady-state LCs (CD1a/CD207^high^). Gates set using isotype controls. Data representative of 8 independent skin donors.
e. Flow cytometry analysis of intranuclear cJUN protein expression in migrated and steady-state LCs (CD1a/CD207^high^). Gates set using isotype controls. Data representative of 2 independent skin donors.
f. Flow cytometry analysis of intranuclear IRF8 protein expression during a timecourse of stimulation with TNF-α in migrated LCs (CD1a/CD207^high^). Gates set using isotype controls. Data representative of 2 independent skin donors.
g. Wild type (wt) and IRF8-/- LCs were isolated from ears and tails from 3 mice/type maintained on C57Bl/6 background. Flow cytometry analysis of intranuclear IRF8 and IRF4 protein expression. n=1, LCs pooled from 3 mice.
h. IFN-γ production by OTI line stimulated with OVA-pulsed wild type (wt) and IRF8-/- LCs. Top panel: live CD3/CD8 OTI cell responses: left: background, right: to H-2Kb restricted OVA257-264 SIINFEKL antigen. Middle panel: live CD3/CD8 OTI cell responses to unpulsed (left) and OVA-pulsed LCs from wt mice. Bottom panel: live CD3/CD8 OTI cell responses to unpulsed (left) and OVA-pulsed LCs from IRF8-/- mice.

**Figure S5. Correlations between antigen processing and cross-presentation signatures and expression of candidate regulatory TF in human primary steady-state LCs transcriptome at a single cell level**

a. Correlation coefficients across DEG involved in antigen cross-presentation in 138 single LCs were compared for candidate regulatory TF. Pearson correlation coefficient is shown for each gene, median ± CI denoted in red.
b. Correlations between the CPTT values for DEGs and IRF4 were computed using corrplot and hmisc packages in R environment (Pearson coefficient >0.11, ^92^). Gene Ontology categories were analysed using ToppGene online Tool. −log (10) for FDR BH p value is shown for the statistically significant categories.

**Figure S6. Coordinated gene expression changes in migrated LCs activated with TNF-α**

a. Transcript-to-transcript clustering, (BioLayout Express3D, r= 0.80; MCL = 1.7) of 1,156 probesets differentially regulated by TNF-α. Lines (edges) represent the similarity between transcripts, circles (nodes) represent genes. DGE: 1156, FDR<0.05, |LogFC|>1. 7 main clusters were identified (gene n>8), denoted by different colours. Enrichment in biological processes was done using ToppGene on-line tool significance denoted by FDR (Benjamini-Hochberg) corrected p-value is shown.
b. Expression profiles in 7 main clusters (gene n>8) identified with transcript-to-transcript clustering, (BioLayout Express3D, r= 0.85; MCL = 1.7) of 1,156 probesets differentially regulated by TNF-α in human migrated LCs.
c. Gene expression of PSME2 CAV1, IRF1 and BATF3 LC assessed by qPCR (expression normalised to house-keeping gene YWHAZ (2-dCT) before (grey bars) and following stimulation with TNF-α (black bars) either during or post-migration. N=2-3 independent skin donors, in duplicate
d. Heat map of genes included in antigen presentation class I signature from Reactime database. Log (2) FPKM median expression values are shown for each gene in the signature. The black line denotes expression cut-off for detection., RNA-seq n=3 donors.
e. Proportion of DEG with increase (M>0) H3K27Ac mark at 2h and 24h in clusters of co-expressed genes up-regulated early (2h, clusters 3) and late (24h, cluster 2) during stimulation with TNF-α. Changes in H3K27Ac acetylation were calculated using MANorm algorithm (MACS2, Zheng 2008) embedded in BioWardrobe tool and genes filtered to include unique common entry across the biological replicates (n=3). Genes with detected changes in acetylation were intersected with DEGs identified by EdgeR analysis
f. UCSC genome browser tracks of H3K27Ac histone mark for B2M from human LC transcriptomic programme. Acetylated area in the promoter region shown by peaks and blue lines along the gene track.
g. UCSC genome browser tracks of H3K27Ac histone mark for PSME2 from human LC transcriptomic programme. Acetylated area in the promoter region shown by peaks and blue lines along the gene track.

### Supplementary Tables

Table S1: Antigen presentation and cross-presentation signatures

Table S2: Correlation of antigen processing signature and TF expression in scRNA Table S3: Genes correlated with IRF4 in scRNA-seq analysis

Table S4: Consensus peaks in H3K4Me3 ChIP-seq in steady-state migrated human LCs Table S5: Consensus peaks in H3K27Ac ChIP-seq in steady-state migrated human LCs

Table S6: GO enrichment in transcript-to-transcript clustering of LC transcriptome over the 24h stimulation with TNF-α

Table S7: Overlaps between chromatin landscape in human migrated LCs and publically available datasets

Table S8: Transcription factor motif enrichment in migrated human LCs

## Acknowledgements

We are grateful to the subjects who participated in this study. We would like to thank Dr Krista Dienger-Stambaugh, Cincinnati Children’s Hospital Medical Center for technical help with processing murine material, and Dr. Nathan Salomonis, for introduction to AltAnalyzer. We acknowledge the use of the IRIDIS High Performance Computing Facility and Flow Cytometry Core Facilities, together with support services at the University of Southampton.

## Author Contributions

MEP and HS: intellectually conceived and wrote the manuscript,

MEP, MAJ, SS, KC, ZW: Biological experiments and flow cytometry, processing of RNA and chromatin

MEP, JW, TW: analysis and meta-analysis of bulk RNA-seq data MEP, JR, MP, XC, MW: analysis of ChIP-seq data

MAJ, PS, MW: reviewing of the manuscript

MEP, PS, MRZ, JW, AV, JD, MacA, SS: processing cells for scRNA-seq, pre-processing and analysis of scRNA-seq data

