## Supplementary Figures for "Genomic programming of antigen cross-presentation in IRF4-expressing human Langerhans cells"

**Supplementary Tables:**

Table S1: Antigen presentation and cross-presentation signatures
Table S2: Correlation of antigen processing signature and TF expression in scRNA
Table S3: Genes correlated with IRF4 in scRNA-seq analysis
Table S4: Consensus peaks in H3K4Me3 ChIP-seq in steady-state migrated human LCs
Table S5: Consensus peaks in H3K27Ac ChIP-seq in steady-state migrated human LCs
Table S6: GO enrichment in transcript-to-transcript clustering of LC transcriptome over the 24h stimulation with TNF-α
Table S7: Overlaps between chromatin landscape in human migrated LCs and publically available datasets
Table S8: Transcription factor motif enrichment in migrated human LCs

**Figure S1**

**a**

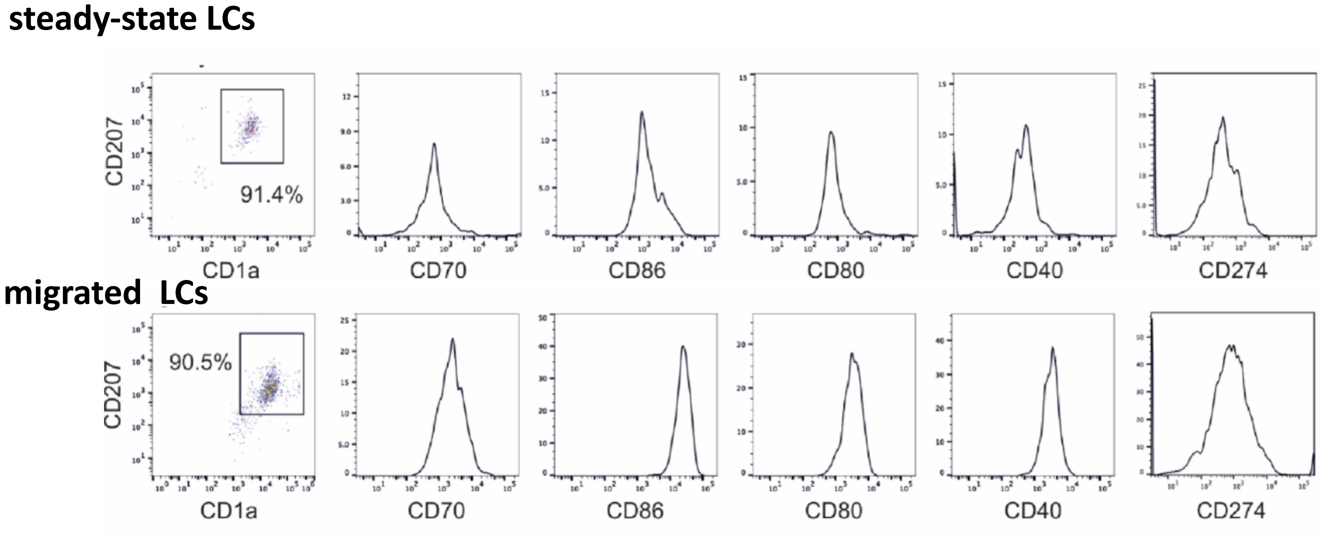
steady-state LCs
CD207
CD1a
91.4%
CD70
CD86
CD80
CD40
CD274
migrated LCs
CD207
CD1a
90.5%
CD70
CD86
CD80
CD40
CD274

**b**

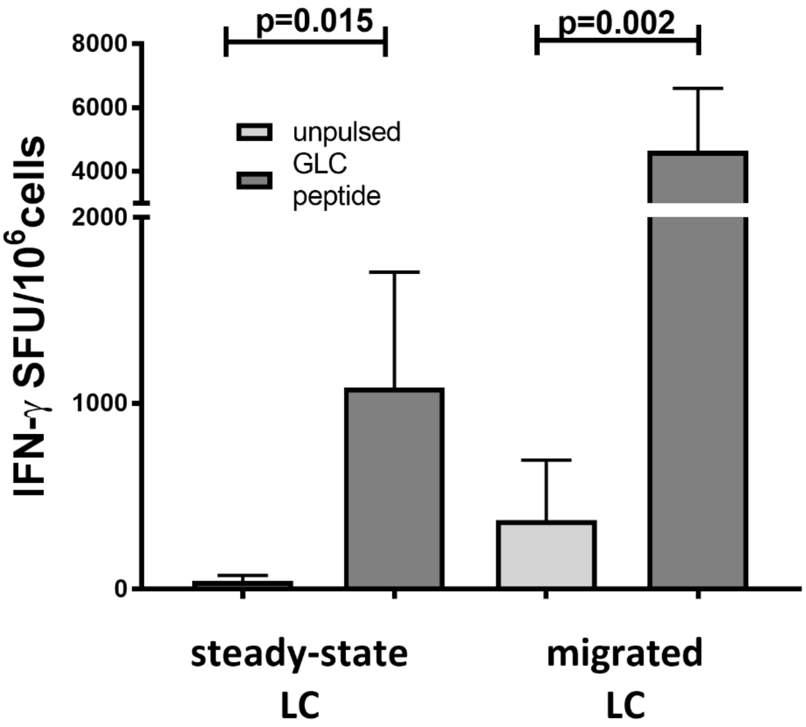
p=0.015
p=0.002
unpulsed
GLC peptide
IFN-γ SFU/10^6 cells
8000
6000
4000
2000
1000
0
steady-state LC
migrated LC

### Figure S2

a

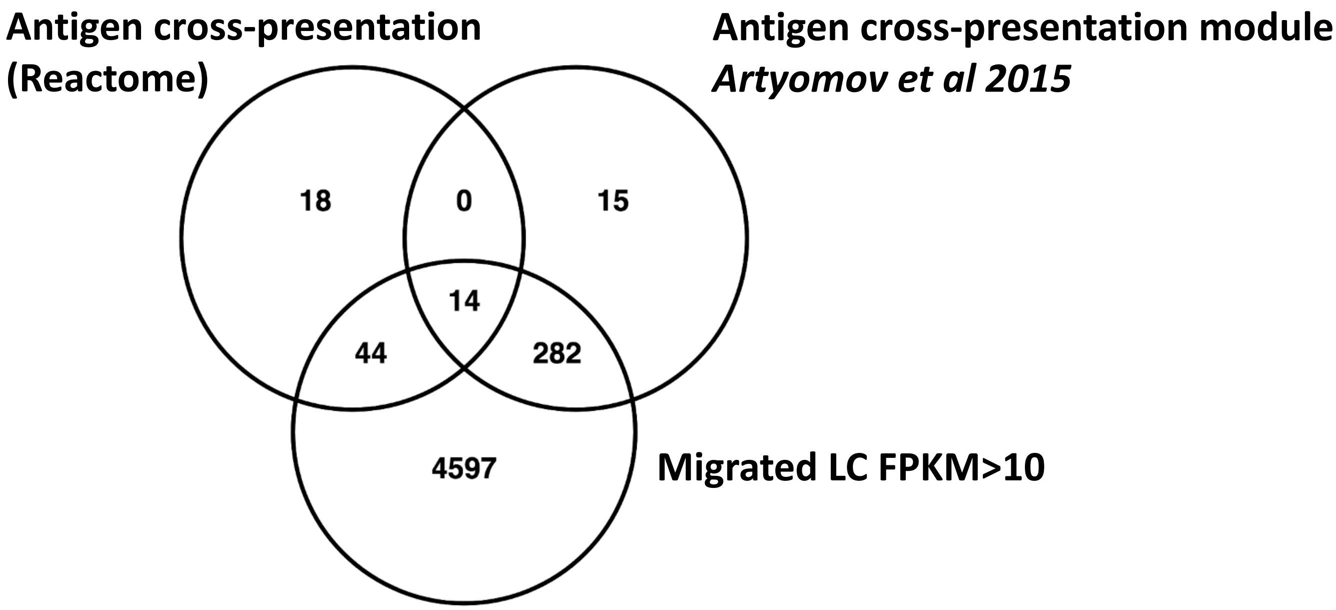
Antigen cross-presentation
(Reactome)
Antigen cross-presentation module
*Artyomov et al 2015*
18
0
15
14
44
282
4597
Migrated LC FPKM>10

b

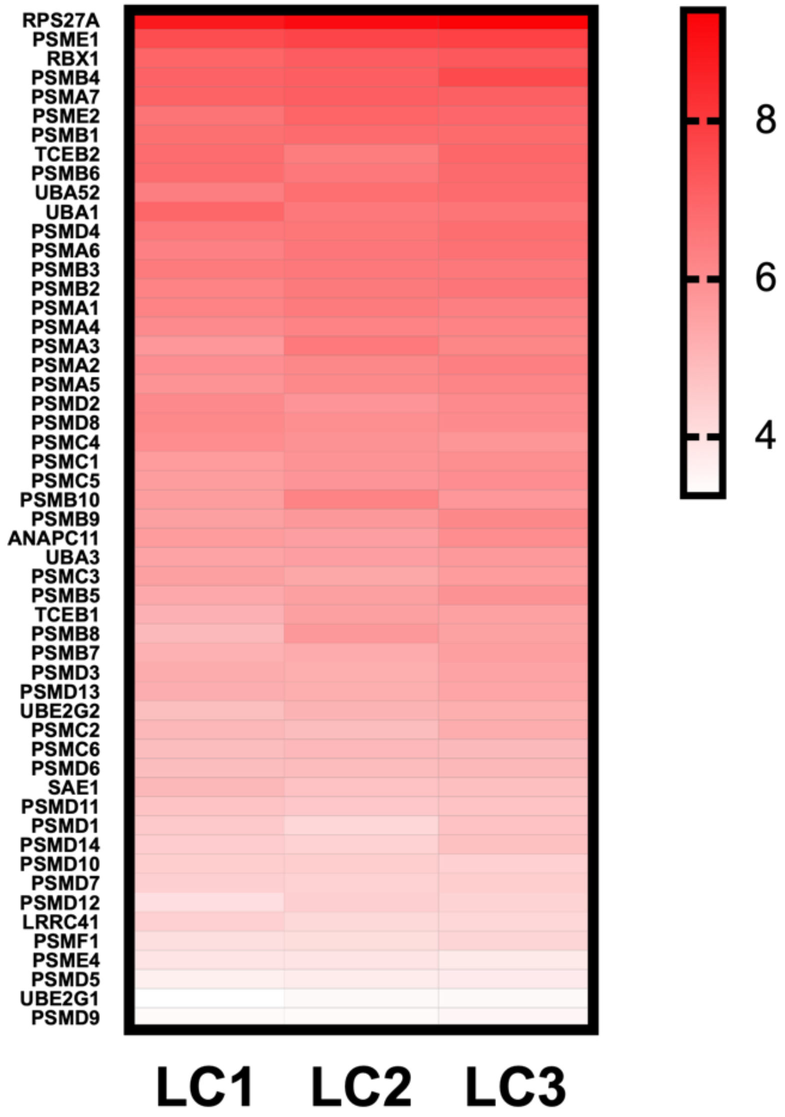
RPS27A
PSME1
RBX1
PSMB4
PSMA7
PSME2
PSMB1
TCEB2
PSMB6
UBA52
UBA1
PSMD4
PSMA6
PSMB3
PSMB2
PSMA1
PSMA4
PSMA3
PSMA2
PSMA5
PSMD2
PSMD8
PSMC4
PSMC1
PSMC5
PSMB10
PSMB9
ANAPC11
UBA3
PSMC3
PSMB5
TCEB1
PSMB8
PSMB7
PSMD3
PSMD13
UBE2G2
PSMC2
PSMC6
PSMD6
SAE1
PSMD11
PSMD1
PSMD14
PSMD10
PSMD7
PSMD12
LRRC41
PSMF1
PSME4
PSMD5
UBE2G1
PSMD9
8
6
4
LC1
LC2
LC3

### Figure S3

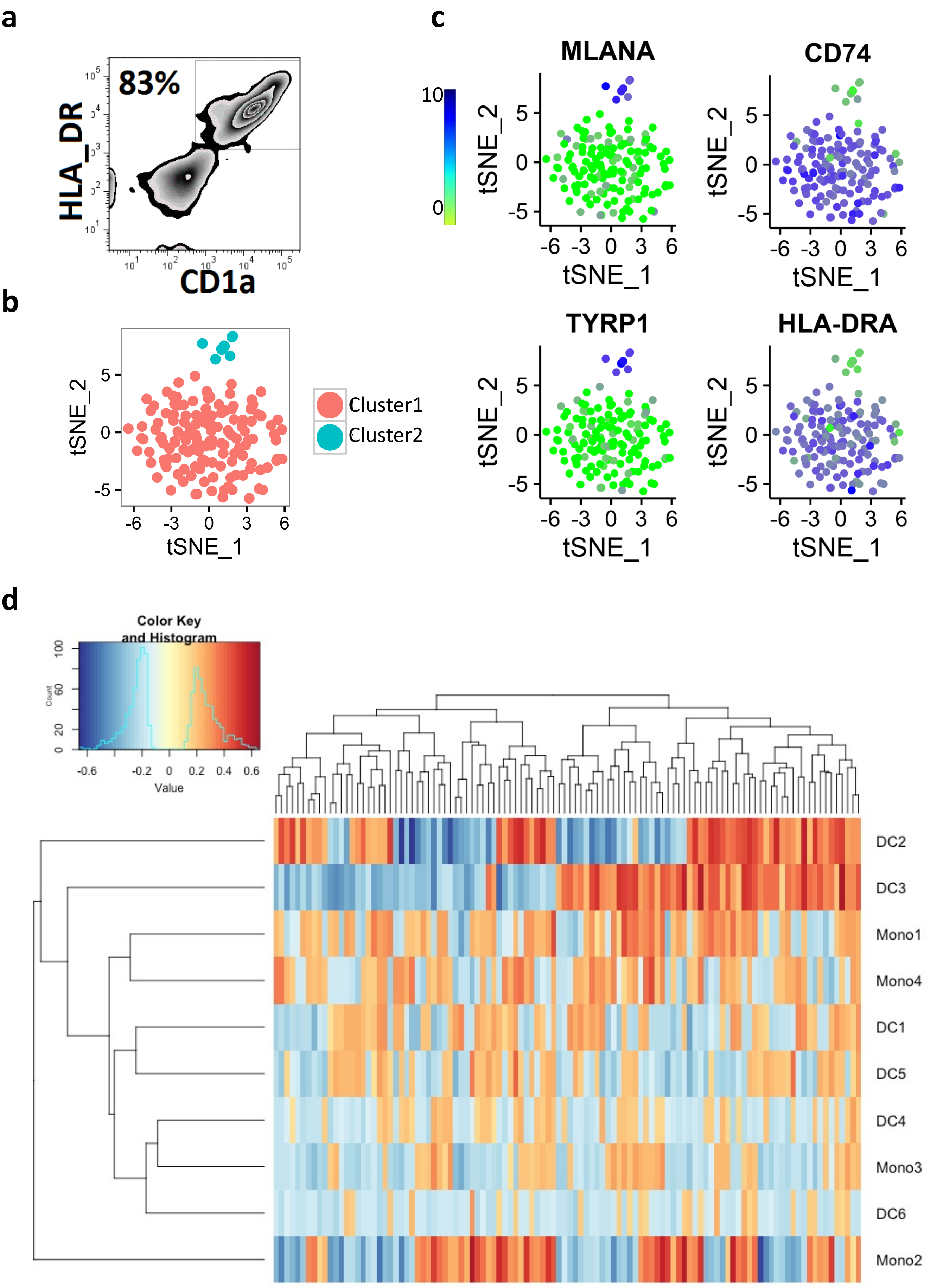

a
83%
HLA_DR
CD1a
b
tSNE_2
tSNE_1
Cluster1
Cluster2
c
MLANA
CD74
TYRP1
HLA-DRA
tSNE_2
tSNE_1
d
Color Key
and Histogram
Count
Value
DC2
DC3
Mono1
Mono4
DC1
DC5
DC4
Mono3
DC6
Mono2

Figure S4

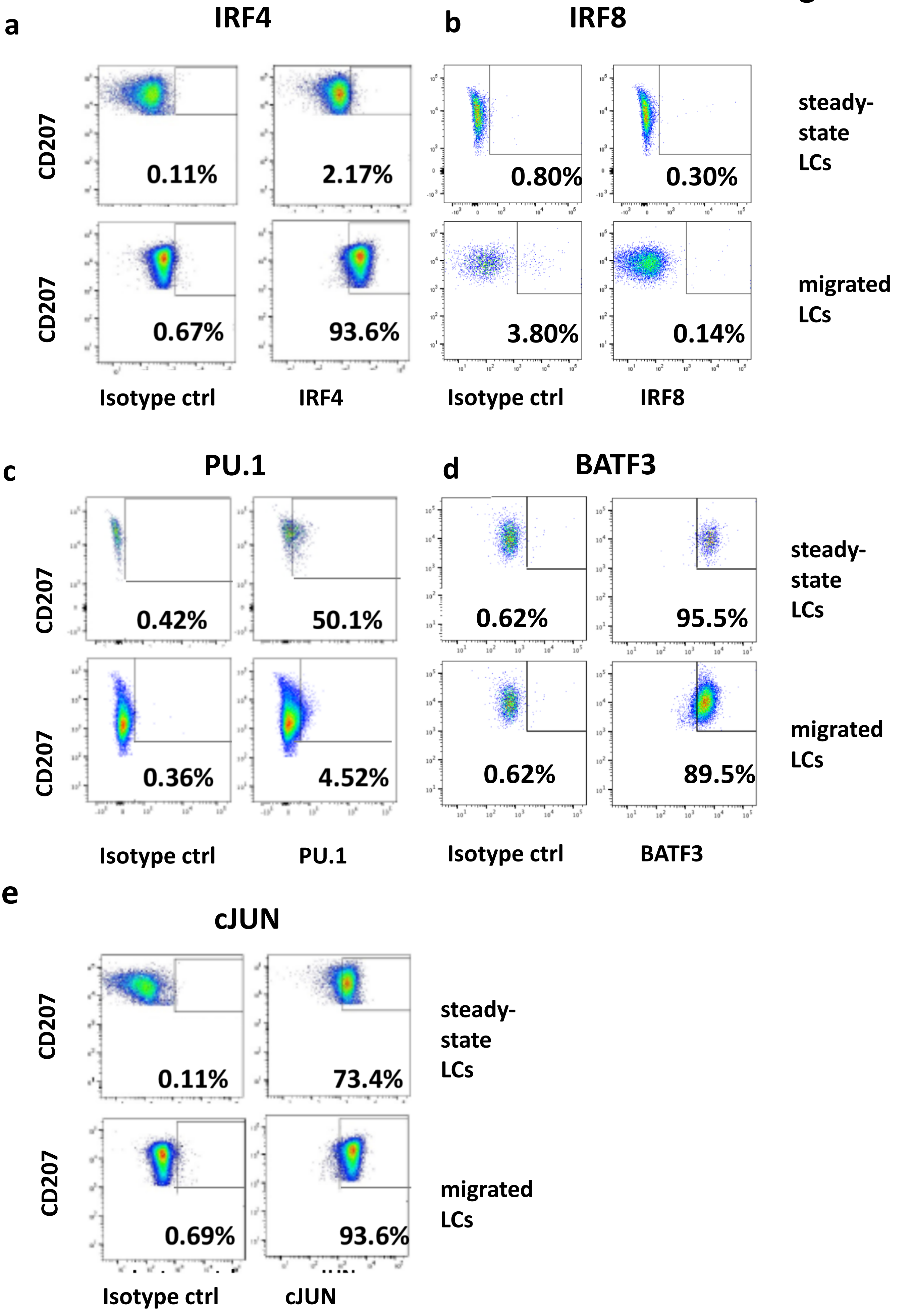
a
IRF4
CD207
0.11%
2.17%
0.67%
93.6%
Isotype ctrl
IRF4
b
IRF8
0.80%
0.30%
3.80%
0.14%
Isotype ctrl
IRF8
steady-state LCs
migrated LCs
c
PU.1
CD207
0.42%
50.1%
0.36%
4.52%
Isotype ctrl
PU.1
d
BATF3
0.62%
95.5%
0.62%
89.5%
Isotype ctrl
BATF3
steady-state LCs
migrated LCs
e
cJUN
CD207
0.11%
73.4%
0.69%
93.6%
Isotype ctrl
cJUN
steady-state LCs
migrated LCs

**f**

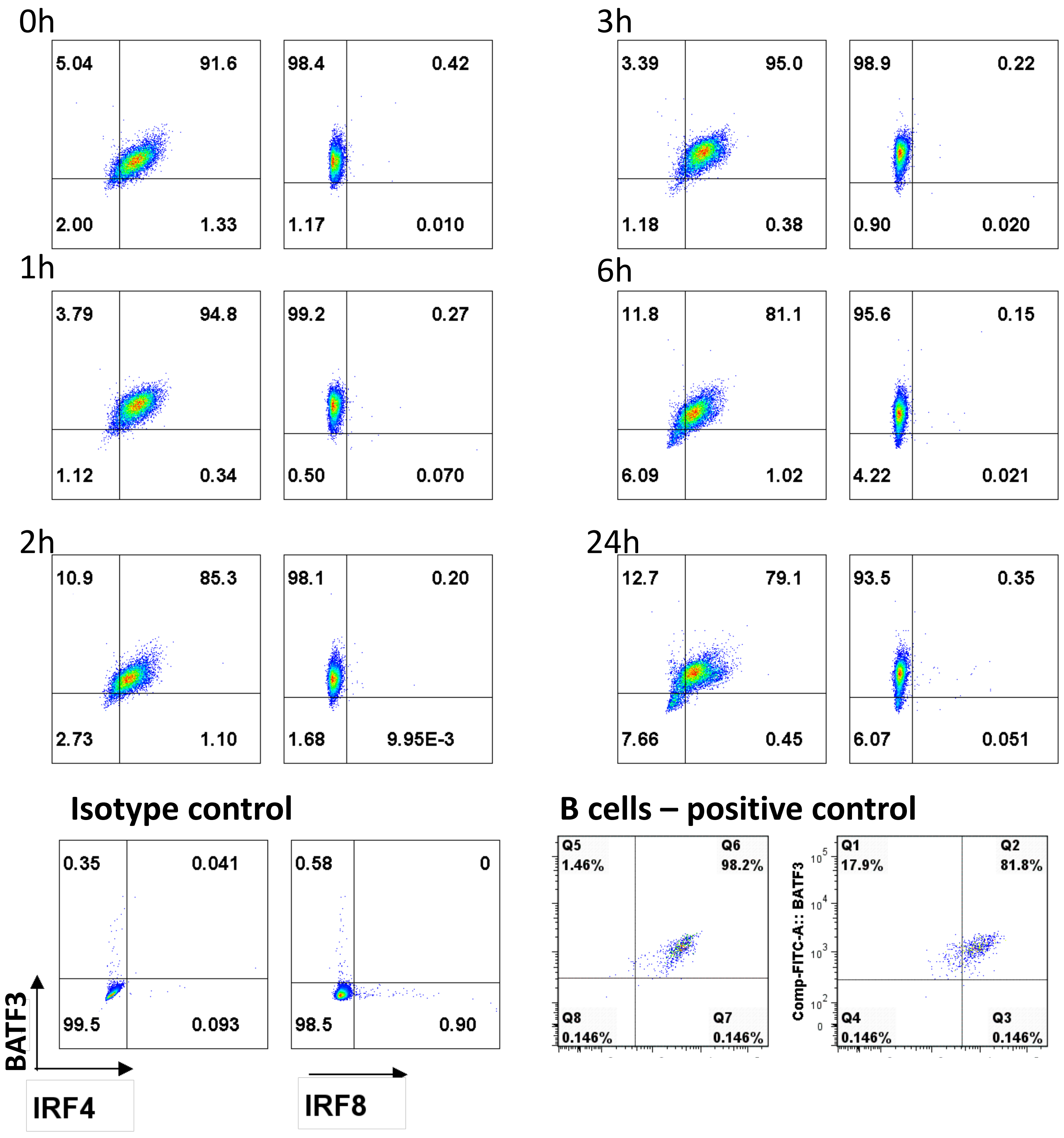

0h
5.04
91.6
2.00
1.33
98.4
0.42
1.17
0.010
1h
3.79
94.8
1.12
0.34
99.2
0.27
0.50
0.070
2h
10.9
85.3
2.73
1.10
98.1
0.20
1.68
9.95E-3
3h
3.39
95.0
1.18
0.38
98.9
0.22
0.90
0.020
6h
11.8
81.1
6.09
1.02
95.6
0.15
4.22
0.021
24h
12.7
79.1
7.66
0.45
93.5
0.35
6.07
0.051
Isotype control
0.35
0.041
99.5
0.093
0.58
0
98.5
0.90
BATF3
IRF4
IRF8
B cells – positive control
Q5 1.46%
Q6 98.2%
Q8 0.146%
Q7 0.146%
Comp-FITC-A:: BATF3
Q1 17.9%
Q2 81.8%
Q4 0.146%
Q3 0.146%

**g**

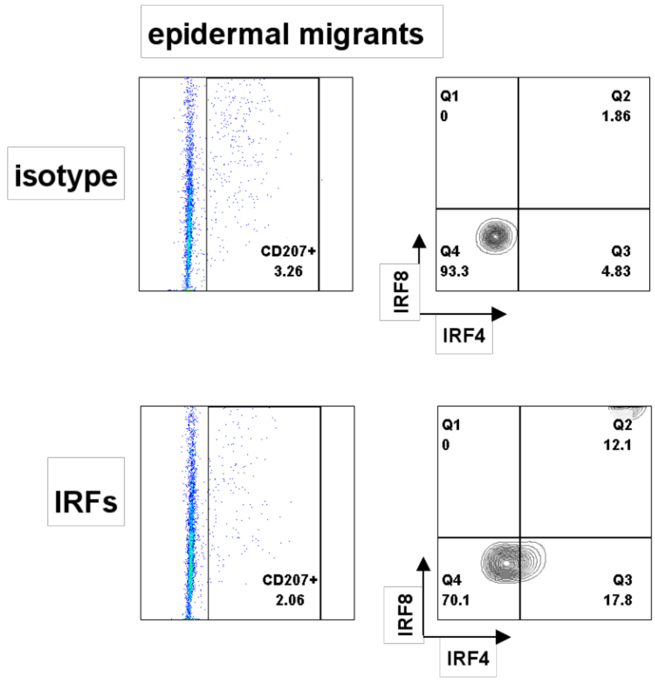
epidermal migrants
isotype
CD207+
3.26
Q1
0
Q2
1.86
Q4
93.3
Q3
4.83
IRF8
IRF4
IRFs
CD207+
2.06
Q1
0
Q2
12.1
Q4
70.1
Q3
17.8
IRF8
IRF4

**h**

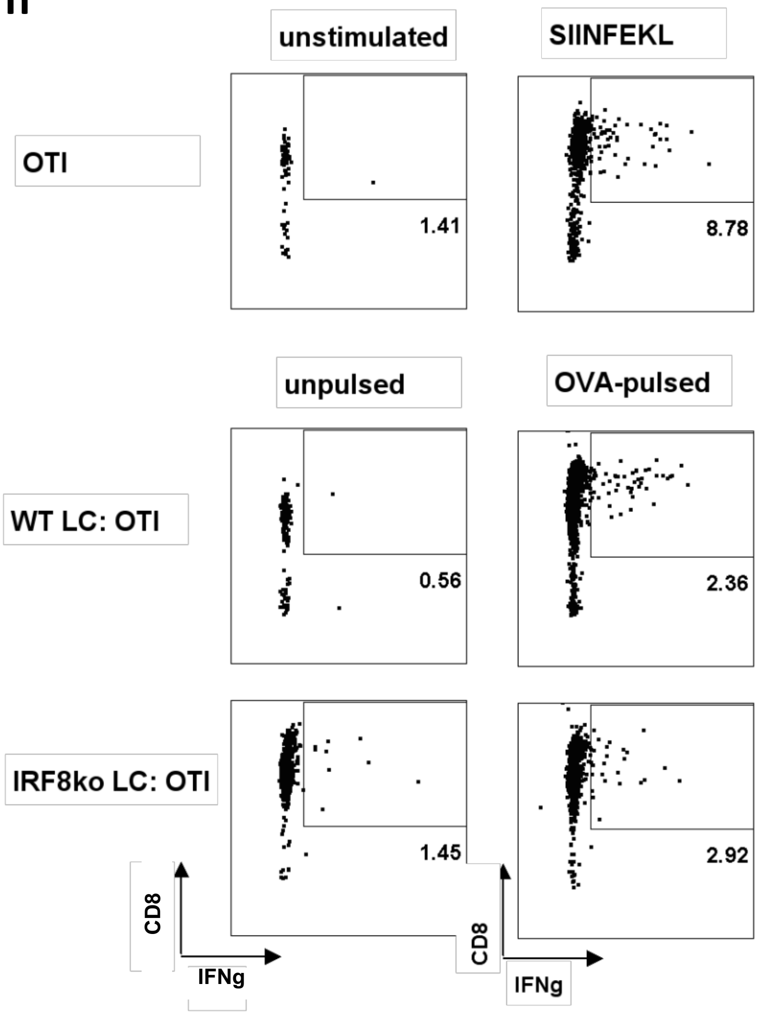
unstimulated
SIINFEKL
OTI
1.41
8.78
unpulsed
OVA-pulsed
WT LC: OTI
0.56
2.36
IRF8ko LC: OTI
1.45
2.92
CD8
IFNg
CD8
IFNg

### Figure S5

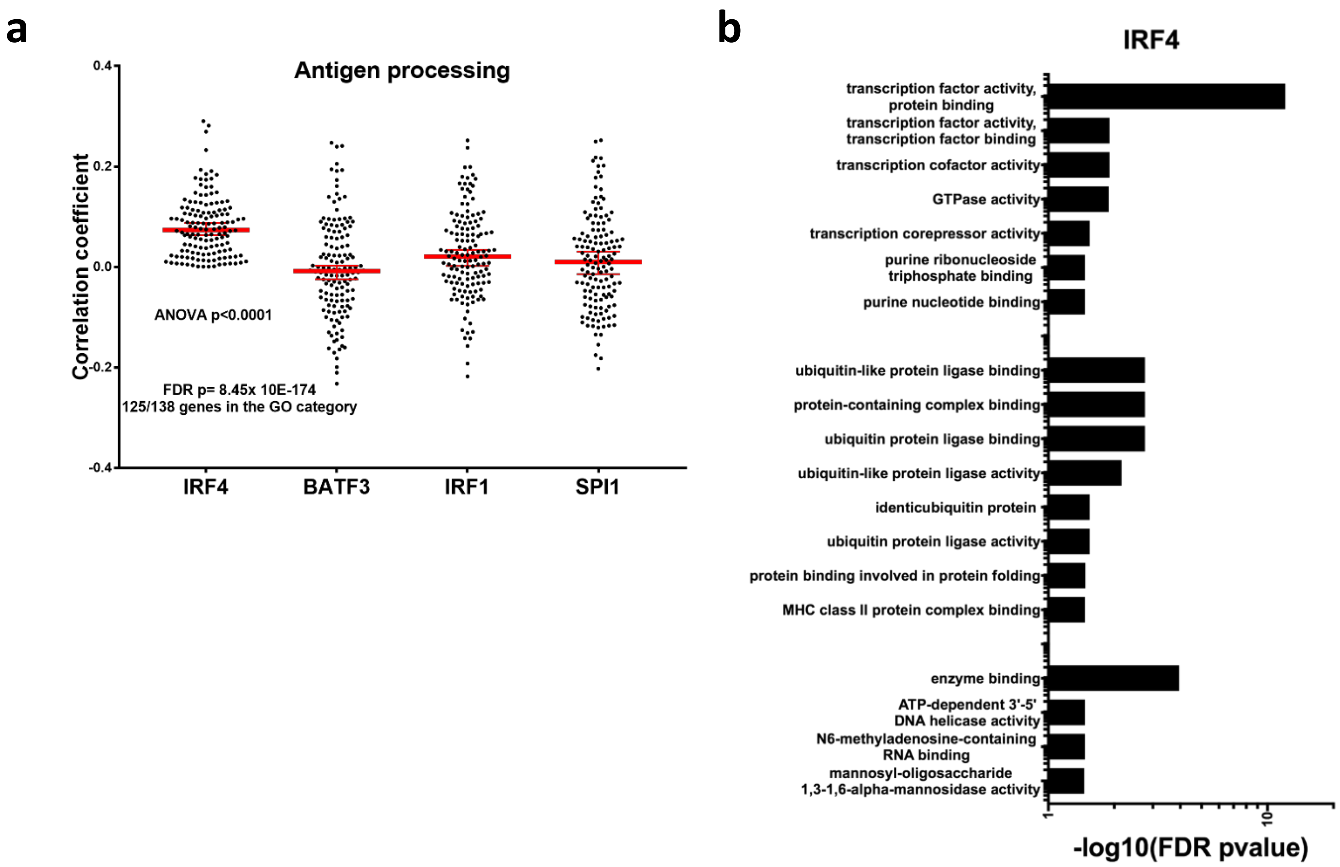
a
Antigen processing
Correlation coefficient
0.4
0.2
0.0
-0.2
-0.4
ANOVA p<0.0001
FDR p= 8.45x 10E-174
125/138 genes in the GO category
IRF4
BATF3
IRF1
SPI1
b
IRF4
transcription factor activity, protein binding
transcription factor activity, transcription factor binding
transcription cofactor activity
GTPase activity
transcription corepressor activity
purine ribonucleoside triphosphate binding
purine nucleotide binding
ubiquitin-like protein ligase binding
protein-containing complex binding
ubiquitin protein ligase binding
ubiquitin-like protein ligase activity
identicubiquitin protein
ubiquitin protein ligase activity
protein binding involved in protein folding
MHC class II protein complex binding
enzyme binding
ATP-dependent 3'-5' DNA helicase activity
N6-methyladenosine-containing RNA binding
mannosyl-oligosaccharide 1,3-1,6-alpha-mannosidase activity
1
10
-log10(FDR pvalue)

Figure S6

a

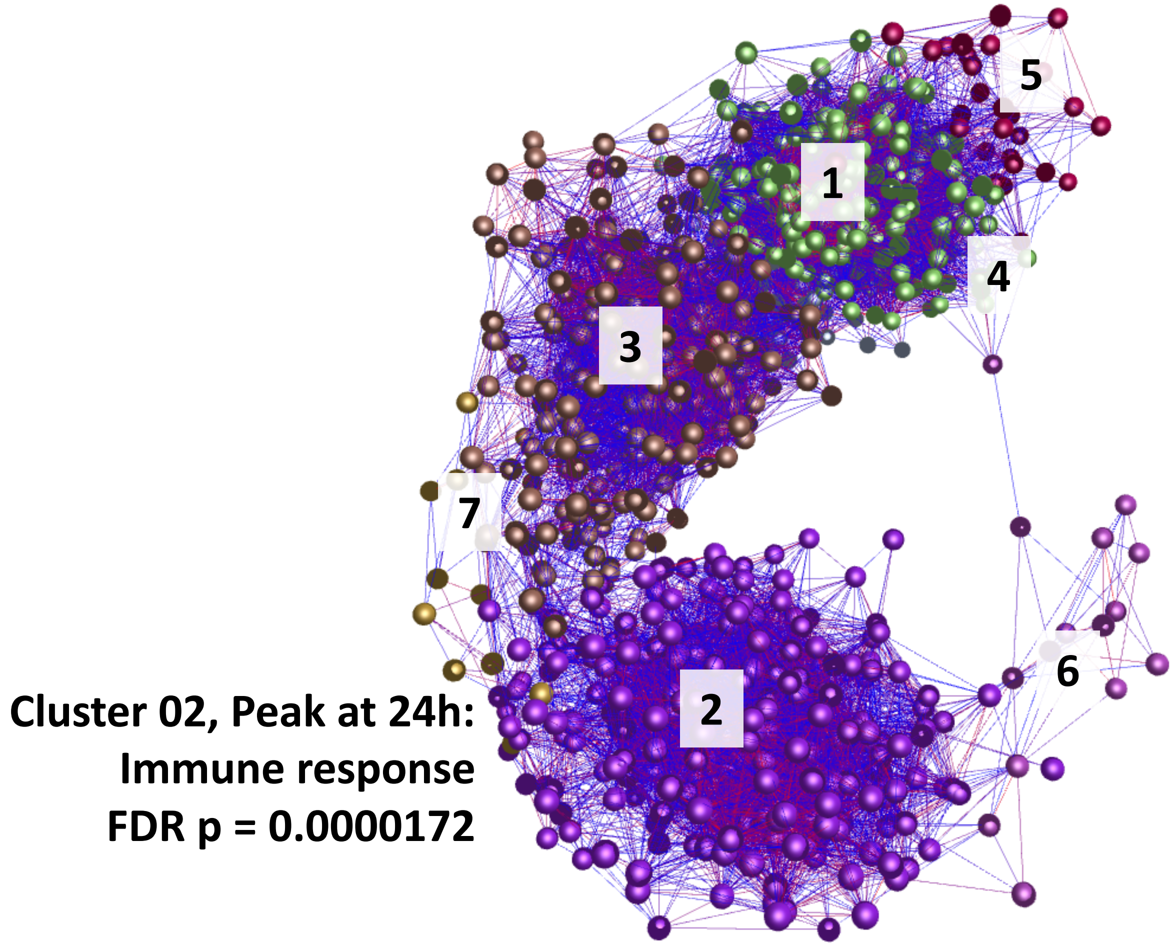
1
2
3
4
5
6
7
Cluster 02, Peak at 24h:
Immune response
FDR p = 0.0000172

b

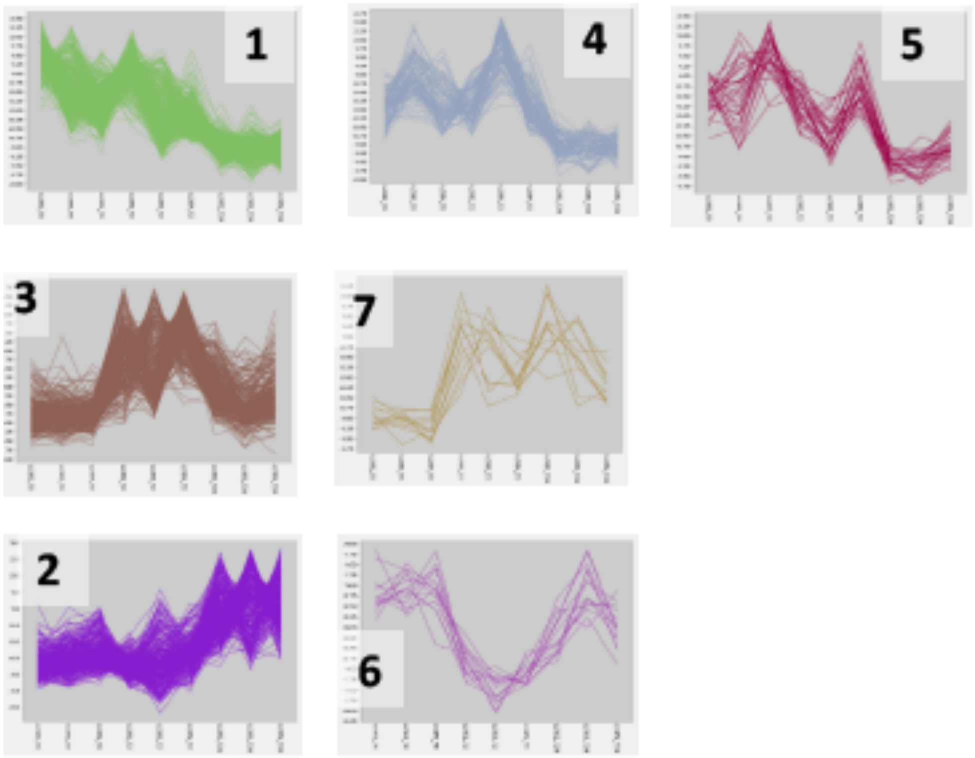
1
4
5
3
7
2
6

**c**

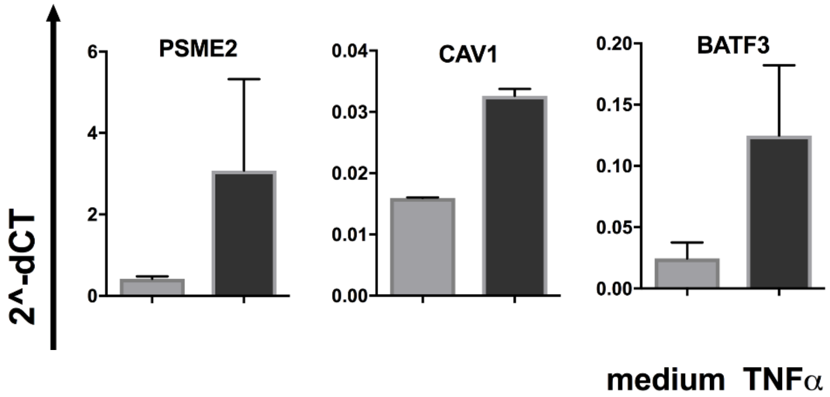
2^-dCT
PSME2
0
2
4
6
CAV1
0.00
0.01
0.02
0.03
0.04
BATF3
0.00
0.05
0.10
0.15
0.20
medium TNFα

**d**

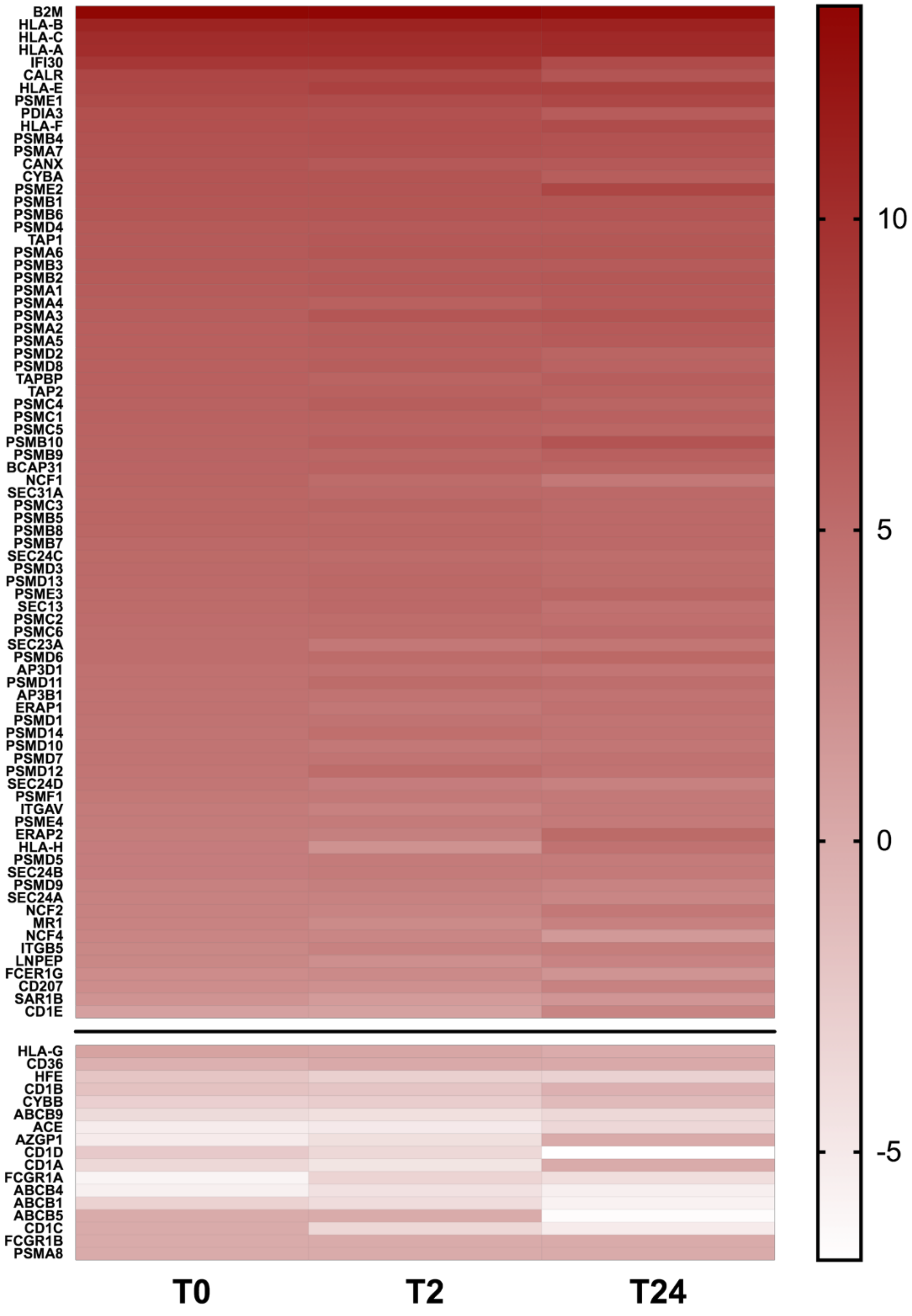
B2M
HLA-B
HLA-C
HLA-A
IFI30
CALR
HLA-E
PSME1
PDIA3
HLA-F
PSMB4
PSMA7
CANX
CYBA
PSME2
PSMB1
PSMB6
PSMD4
TAP1
PSMA6
PSMB3
PSMB2
PSMA1
PSMA4
PSMA3
PSMA2
PSMA5
PSMD2
PSMD8
TAPBP
TAP2
PSMC4
PSMC1
PSMC5
PSMB10
PSMB9
BCAP31
NCF1
SEC31A
PSMC3
PSMB5
PSMB8
PSMB7
SEC24C
PSMD3
PSMD13
PSME3
SEC13
PSMC2
PSMC6
SEC23A
PSMD6
AP3D1
PSMD11
AP3B1
ERAP1
PSMD1
PSMD14
PSMD10
PSMD7
PSMD12
SEC24D
PSMF1
ITGAV
PSME4
ERAP2
HLA-H
PSMD5
SEC24B
PSMD9
SEC24A
NCF2
MR1
NCF4
ITGB5
LNPEP
FCER1G
CD207
SAR1B
CD1E
HLA-G
CD36
HFE
CD1B
CYBB
ABCB9
ACE
AZGP1
CD1D
CD1A
FCGR1A
ABCB4
ABCB1
ABCB5
CD1C
FCGR1B
PSMA8
T0
T2
T24
10
5
0
-5

e

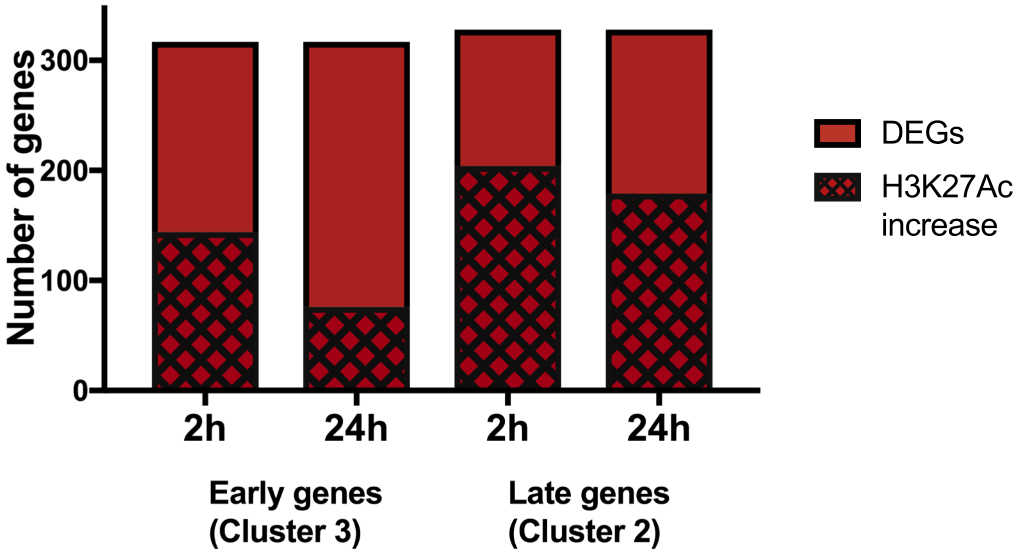


f

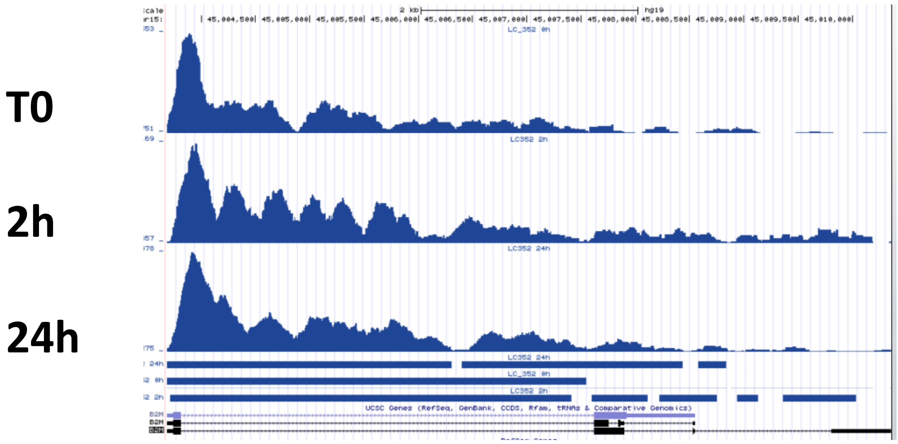


H3K27Ac B2M

g

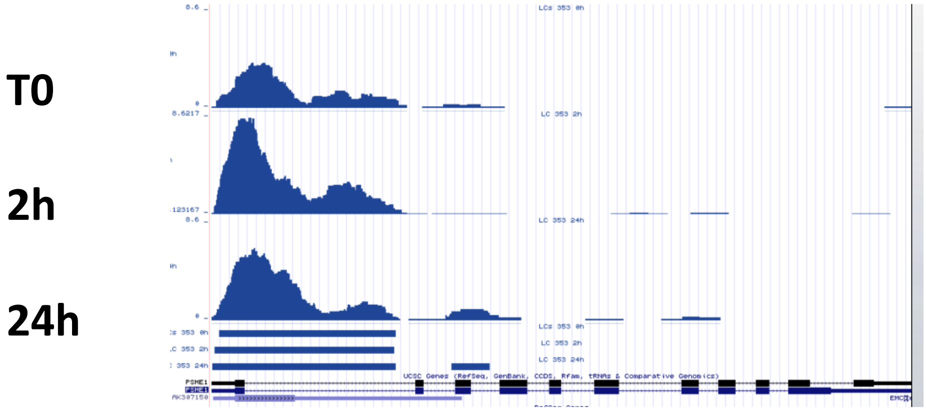


H3K27Ac PSME2

h

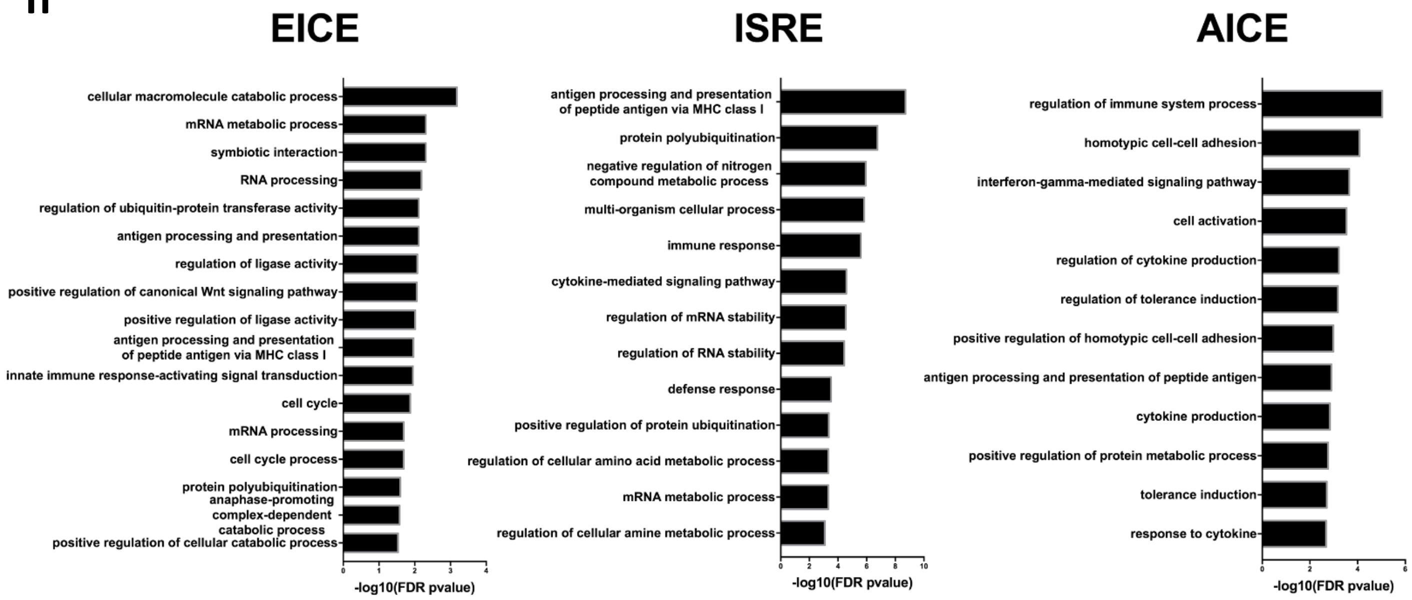

EICE
cellular macromolecule catabolic process
mRNA metabolic process
symbiotic interaction
RNA processing
regulation of ubiquitin-protein transferase activity
antigen processing and presentation
regulation of ligase activity
positive regulation of canonical Wnt signaling pathway
positive regulation of ligase activity
antigen processing and presentation of peptide antigen via MHC class I
innate immune response-activating signal transduction
cell cycle
mRNA processing
cell cycle process
protein polyubiquitination
anaphase-promoting complex-dependent catabolic process
positive regulation of cellular catabolic process
-log10(FDR pvalue)
ISRE
antigen processing and presentation of peptide antigen via MHC class I
protein polyubiquitination
negative regulation of nitrogen compound metabolic process
multi-organism cellular process
immune response
cytokine-mediated signaling pathway
regulation of mRNA stability
regulation of RNA stability
defense response
positive regulation of protein ubiquitination
regulation of cellular amino acid metabolic process
mRNA metabolic process
regulation of cellular amine metabolic process
-log10(FDR pvalue)
AICE
regulation of immune system process
homotypic cell-cell adhesion
interferon-gamma-mediated signaling pathway
cell activation
regulation of cytokine production
regulation of tolerance induction
positive regulation of homotypic cell-cell adhesion
antigen processing and presentation of peptide antigen
cytokine production
positive regulation of protein metabolic process
tolerance induction
response to cytokine
-log10(FDR pvalue)
